# Human macula formation involves two waves of retinoic acid suppression via *CYP26A1* that modulate cell cycle exit and cone subtype specification

**DOI:** 10.1101/2024.09.18.613197

**Authors:** Philippa Harding, Maja Wojtynska, Alexander J. Smith, Robin R. Ali, Rachael A. Pearson

## Abstract

The human macula is a specialized, cone-rich region of the eye, critical for high-acuity vision, yet the pathways regulating its development remain poorly understood. RA-catabolizing enzyme *CYP26A1* establishes the chick high-acuity area via upregulation of *FGF8*. However, detailed analysis of this pathway and its functions has not been performed in early human fetal tissue. Fluorescent *in situ* hybridization revealed striking biphasic *CYP26A1* expression but little *FGF8* in the presumptive macula region between post-conception weeks (PCW)6-17. Pharmacological RA signaling inhibition in human retinal organoids mimicking the two waves of *CYP26A1* revealed early RA inhibition prompted early cell cycle exit and increased cone genesis, while late inhibition altered cone subtype specification. Conversely, recombinant FGF8 had no effect on photoreceptor fate. This work provides spatiotemporal examination of *CYP26A1* across human macular development, as well as experimental evidence for the different roles of RA signaling inhibition in a human model of retinal development.

## Introduction

The macula is a 5-6mm region at the center of the retina, containing within it the fovea, an area critical for high-acuity and color vision in humans and simian primates(Hoshino *et al*., 2017; Voigt *et al*., 2021). Damage and degeneration in this region lead to debilitating sight loss, significantly impacting activities such as reading and recognizing faces. The macula has a unique architecture comprising a high density of cone photoreceptors, contrasting with the rod-dominant peripheral retina, the highest density (200,000 cones/mm^2^) being at the center of the macula in the foveola, a 300μm avascular rod-free zone (RFZ) consisting exclusively of medium (M/green) and long (L/red) wavelength-detecting cones(Curcio *et al*., 1990; Voigt *et al*., 2021). The proportion of cones:rods declines with increasing eccentricity, being ∼1:1 in the fovea, dropping to 1:30 in the peripheral retina. Additionally, the macula contains the highest density of retinal ganglion cells (RGCs) in the retina, and a highly ordered synaptic configuration to aid rapid phototransduction.

The retina exhibits a striking center-to-periphery gradient of development, the central retina entering neurogenesis ∼50 days before peripheral regions(Hoshino *et al*., 2017), whilst macular differentiation precedes even the equivalent region in the nasal retina, with immature cones present in the presumptive macula (PM) by the end of post-conception week (PCW)8/Carnegie stage (CS)23(Hendrickson and Zhang, 2019). However, due to central migration of cones into the fovea, adult cone density is not achieved until 4-6 years(Curcio *et al*., 1990). Despite their importance, the molecular mechanisms underlying human macular and foveal development remain poorly understood, owing to the lack of equivalent structures in classical animal models. Furthermore, minimal evaluation of macular formation has been performed in human tissue prior to PCW8, due to limited tissue samples, and the lack of a macular marker expressed at this early stage(Hendrickson, 2016). Some other species do possess a broadly analogous high-acuity region; Chickens possess an RFZ, called the *area centralis* or High-Acuity Area (HAA)(Mey and Thanos, 2000). Recent research has shown that chick HAA formation is dependent on regulation of retinoic acid (RA) levels(Silva and Cepko, 2017). RA is a morphogen that can control gene expression through binding to RA receptors (RARs) and retinoid X receptors (RXRs), which then engage RA-response elements (RAREs)(Cvekl and Wang, 2009). Upon binding to RAREs, co-activators are recruited and lead to transcriptional regulation of specific RA-regulated genes(Cunningham and Duester, 2015). RA is synthesized by retinal dehydrogenases (ALDHA1-3) and catabolized by cytochrome P450 family 26 enzymes (CYP26s) (Cvekl and Wang, 2009).

In the developing chick retina, strong expression of *Cyp26a1/c1* was observed at a single spot at the presumptive HAA early, coincident and colocalized with similarly strong expression of *Fgf8* (fibroblast growth factor 8), which is otherwise downregulated by RA(Cunningham and Duester, 2015; Silva and Cepko, 2017). Accordingly, transient addition of RA led to loss of *Fgf8* expression, and disruption of the RFZ and a decrease in the high RGC density typical of the HAA(Silva and Cepko, 2017). In the same study, expression of *CYP26A1* mRNA was observed in the human PM at CS22 (late PCW8), indicating a potentially conserved signaling pathway in HAA/macula development. In support of this, increased expression of *Cyp26a1* by Müller glia (MG) has been observed, albeit much later in development, in the zebrafish HAA(Lahne, Yoshimatsu and MacDonald, 2023), macaque and marmoset(Peng *et al*., 2019; Krueger *et al*., 2024), and bulk and scRNAseq studies have reported *CYP26A1* to be significantly upregulated in human macula compared with peripheral retina in mid-gestation (PCW20) and in the MG of adult retina human retina(Peng *et al*., 2019; Cowan *et al*., 2020). Notably, recent detailed analysis of *CYP26A1* and *FGF8* expression in macaque(Krueger *et al*., 2024) found that *CYP26A1*, but not *FGF8*, is highly expressed in the developing macular region at day 40, after the onset of cone genesis (corresponding to human ∼PCW10(Hendrickson and Zhang, 2019)). However, little is known about the spatiotemporal expression of *CYP26A1* and *FGF8* in human fetal retina across the whole period of macular development. Moreover, while various roles have been ascribed to RA signaling across retinal development, the precise downstream functions of RA inhibition in human macular formation are not known.

Stem cell-derived human stem cell-derived retinal organoids (hROs) broadly resemble the human neural retina and closely follow the timing of human retinal development(Gonzalez-Cordero *et al*., 2017). As such, they provide an opportunity to explore the signaling pathways that regulate human retinal development. Here, we used *in situ* hybridization in human fetal tissue and pharmacological manipulation in hROs to investigate the spatiotemporal expression and potential roles of RA and *FGF8* in human macular development.

## Results

### The RA catabolizing enzyme *CYP26A1* is expressed in two distinct waves within the developing human macula

To determine the spatiotemporal expression of *CYP26A1* and *FGF8,* we performed quantitative RNAscope™ *in situ* hybridization on human fetal retina samples between PCW6 (CS16-17) and PCW17 (**Fig.1**). No distinct labelling for *CYP26A1* was visible anywhere in the retina at PCW6, but at PCW7 (CS18-19) weak, specific expression was just visible and localized to a single spot approximately 1mm temporal to the optic nerve head (ONH) (**Fig.1a**), a location identified previously as the PM(Hendrickson, 2016). Within this region, *CYP26A1* was distributed throughout the neuroblastic layer (NBL), colocalizing with VSX2+ retinal progenitor cells (RPCs) (**Fig.1bi**).

**Fig. 1.**
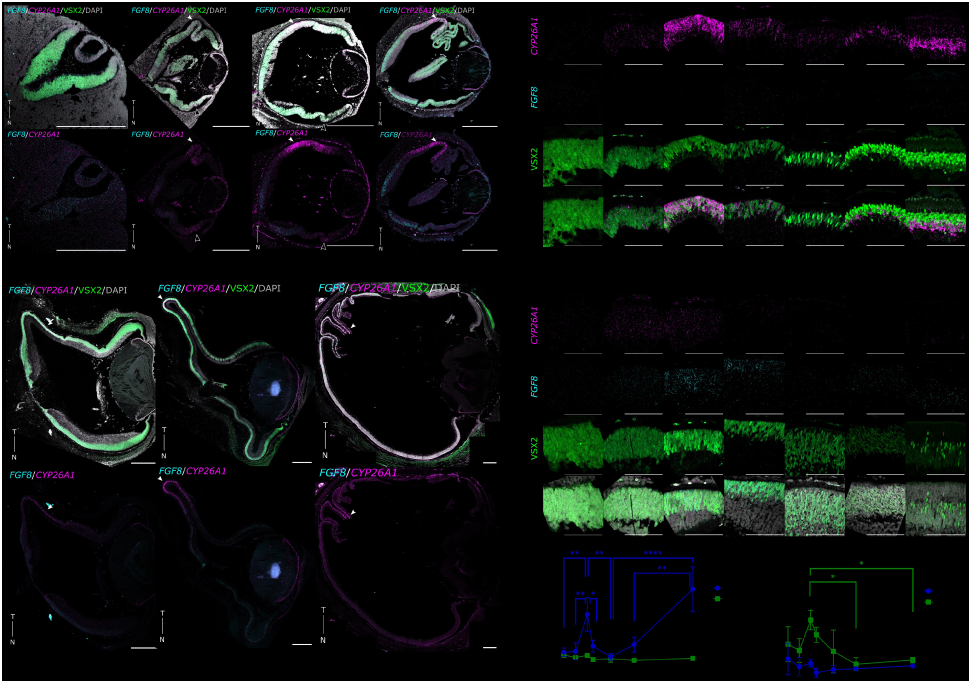
*CYP26A1* shows focal, biphasic expression within the PM of PCW6-17 human fetal retina, peaking at early PCW8 and PCW16/17. (a) Human fetal retina samples from PCW6-17 stained for *FGF8 (*cyan) and RA catabolizing enzyme *CYP26A1* (magenta), showing *CYP26A1*+ spot-like PM region colocalizing with RPC (retinal progenitor cell) marker, VSX2 (green) in the neuroblastic layer between PCW7-8 (CS18-23), and basal to the VSX2+ layer at PCW12-17. (b) Zoom-ins of the (i) PM and (ii) ONH, (c) Quantification of *CYP26A1*/*FGF8* fluorescence intensity over PCW6-17 in PM/ONH, normalized to positive control (*POLR2A*/*PPIB*). One-way ANOVA yielded a statistically significant difference in *CYP26A1* expression in the PM between at least 2 timepoints (*F*(6,15)=13.96, *p*<0.0001), and Tukey’s test found significant increase in mean fluorescence intensity between PCW7-early PCW8 (CS20-21) (*q*(15)=6.34, *p*=0.006), and a significant decrease between early PCW8-late PCW8 (CS22-23) (*q*(15)=5.90, *p*=0.01). ANOVA found statistical differences in *CYP26A1* expression in the ONH (*F*(6,14)=3.39,*p*=0.03), however, no significance was identified by post-hoc tests. ANOVA yielded no significant differences in *FGF8* expression in the PM (*F*(6,15)=1.18,*p*=0.40) between any time points. Conversely, statistical differences were found in *FGF8* in the ONH (*F*(6,15)=4.06,*p*=0.01), with significantly higher *FGF8* at PCW8 compared to PCW12 (*q*(15)=5.45, *p*=0.02) and PCW17 (*q*(15)=4.95, *p*=0.04). Images of probe labelling through entire CS20 retina, positive and negative controls, intensity measurements and all multiple comparison results for PM *CYP26A1* expression are shown in **Fig.S1**. White arrows indicate PM, while white outlined arrow heads indicate low levels of *CYP26A1* expression visible at nasal edge of the retina. Sections counterstained with DAPI (greyscale). T: Temporal; N: Nasal; CS: Carnegie Stage; PCW: Post-conception Weeks, PM: Presumptive macula. \**p* value<0.05, \*\**p* value<0.01, *** *p* value<0.001. Data shown as mean±SD. Images taken at (a) 20x/(b) 40x, representative of n≥2 samples/timepoint. Scale bars: (a) 500μm, (b) 100μm.

At the start of PCW8 (CS20-21, herein termed “early PCW8”), significantly higher levels of *CYP26A1* were observed in the PM within the VSX2+ NBL, compared to PCW7 (*q*(15)=4.79, *p*=0.05; **Fig.1a,bi,ci**), with expression significantly reducing again by the end of PCW8 (CS22-23, herein termed “late PCW8”; *q*(15)=6.34, *p*=0.006; **Fig.1a,bi,ci**). *CYP26A1* was also expressed in the lens epithelium (LE) and at low levels in the nasal edge of the retina and around the ONH at PCW7-early PCW8 (**Fig.1a; Fig.S1a**). To provide context regarding retinal maturation, at late PCW8 RPCs expressing the cone/bipolar progenitor marker OTX2 were distributed sparsely throughout the NBL of the entire retina, but were more densely concentrated in the PM, where they were located towards the apical side of the retina (**Fig.2ai-iv**). The RXRγ+ nascent GCL was visible throughout the retina, but only a few RECOVERIN+/RXRγ+ cone photoreceptor cells were visible, in the PM (**Fig.2ai-ii**).

By PCW10, *CYP26A1* mRNA expression was significantly reduced in the PM region, compared to early PCW8 (*q*(15)=7.28, *p*=0.002; **Fig.1a,bi,ci**); moreover, its expression was restricted to a band of VSX2+/OTX2+ cells located in the nascent inner nuclear layer (INL) (**Fig.2bi-iv**), At PCW10, RXRγ+/RECOVERIN+/OTX2+ cone photoreceptors were located exclusively within a single cell layer (the nascent outer nuclear layer; ONL) at the apical edge of the PM, which did not express *CYP26A1* (**Fig. 2bi- iv**). RXRγ+/RECOVERIN+ cones were not present in the peripheral retina at this stage (**Fig. 2bi**).

At PCW12, *CYP26A1* expression was visibly increased within the PM, compared with PCW10, localizing to the basal edge of the VSX2+ INL, although this was not statistically significant (**Fig.1a,bi,ci**). However, by PCW17, expression of *CYP26A1* was significantly higher than PCW10 (*q*(15)=9.71, *p*<0.0001; **Fig.1a,bi,ci)** and PCW12 (*q*(15)=7.23, *p*=0.002; **Fig.1a,bi,ci; Fig.S1c**). At this stage, *CYP26A1* expression was exclusively within the PM (**Fig.1a**) and almost entirely restricted to VSX2+/SOX9+ MG cells, with minimal signal visible in the VSX2+/SOX9- bipolar cells or the ONL (**Fig.2c**).

**Fig. 2.**
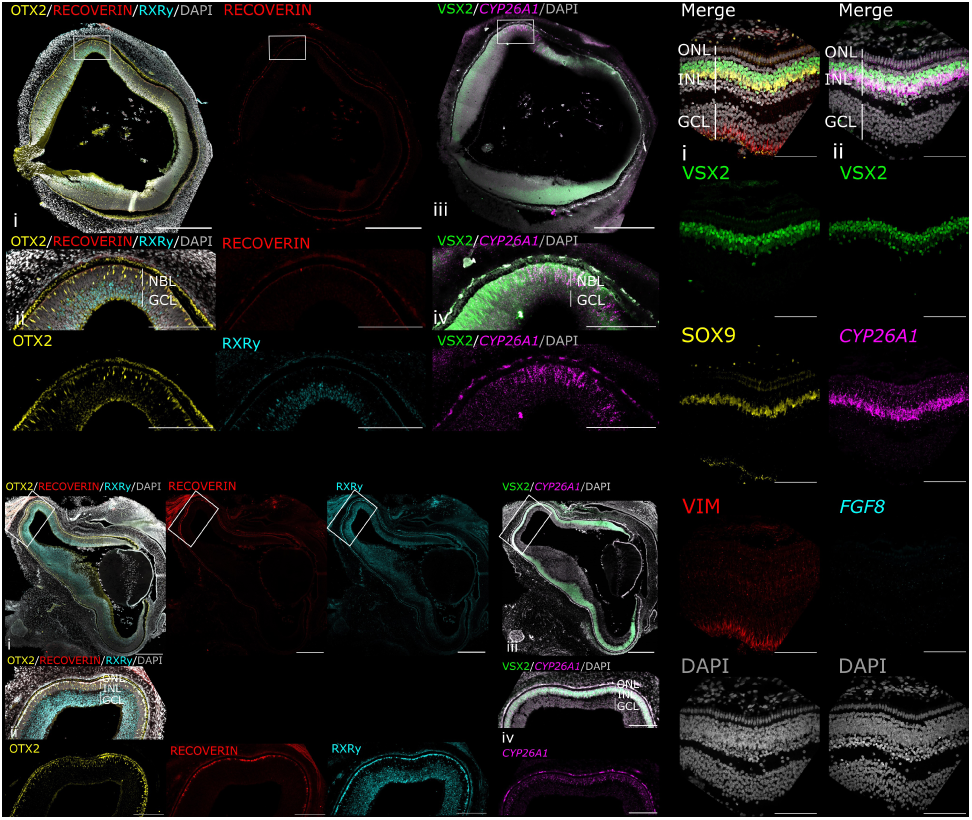
Cone progenitor markers are expressed in the NBL of late PCW8 PM, localized to the INL by PCW10 but do not co-express *CYP26A1*, while macular Müller Glia (MG) cells express *CYP26A1* in PCW17 human fetal retina. (a), At late PCW8 (CS23), OTX2 (yellow) is expressed in cone/bipolar-biased RPCs in the neuroblastic layer (NBL) throughout the retina, at highest density in the PM, while no RECOVERIN (red) or RXRγ+ (cyan) cone precursor markers are visible. (b), At PCW10, the PM region expresses a thick layer of OTX2+ cells in the INL, and RXRγ+/RECOVERIN+ cone precursors localize to the apical single-cell thick ONL layer of the central retina, not present in the peripheral retina. *CYP26A1* expression is restricted to the INL, not expressed in the RECOVERIN+/RXRγ+/OTX2+ cone precursors in the ONL. (c) At PCW17, *CYP26A1* (magenta) is visible in the PM, localized to the VSX2+ (green)/SOX9+ (yellow) developing Müller glia layer, basal to the VSX2+/SOX9- developing bipolar cell layer. Müller glia cytoskeletal marker Vimentin (VIM, red) is clearly expressed, while no *FGF8* (cyan) is visible in the PM. Stains were performed on consecutive sections from the same respective retina for ai/iii, bi/iii and ci/ii, and counterstained with DAPI (greyscale). CS: Carnegie Stage; PCW: Post-conception Weeks; NBL: Neuroblastic Layer; GCL: Ganglion Cell Layer; ONL: Outer Nuclear Layer; INL: Inner Nuclear Layer. Images taken at (ai/bi) 20x/(aii-iii/bii-iii/c) 40x. White box indicates magnified region of interest. Images representative of n≥2 samples. Scale bars: (a) 500μm, (b-c) 100μm.

### Spatiotemporal *FGF8* expression pattern is not correlated with the human PM

In the chick, *Fgf8* is initially expressed in a comet-like pattern, with a broad stripe of lower expression extending temporally and a strong, central spot of expression overlapping with the spot-like expression of *Cyp26a1/c1* within the HAA(Silva and Cepko, 2017). In human fetal retina, positive *FGF8* mRNA labelling was visible around the ONH throughout PCW6-17 (**Fig.1a, Fig.S1a-b**), with the highest expression seen at PCW8 (**Fig.1bii,cii**; **Fig.3**). Examining the whole retina, expression exhibited a strong ONH-to-periphery gradient (**Fig.1a, Fig.3a, Fig.S1b**), similar to the developing macaque (d40)(Krueger *et al*., 2024), mouse (E9.5-14.5)(Crossley and Martin, 1995),and chick eye (E2-E7)(Soukkarieh *et al*., 2007), and from PCW12, *FGF8* labelling was also observed in the very peripheral retina, near the lens (**Fig.1a**). However, in contrast to the clearly demarcated, spot-like and spatiotemporally coincident expression of *Fgf8* and *Cyp26a1/c1* seen in the HAA of the chick, very little *FGF8* expression was observed within the *CYP26A1*-positive PM (**Fig.1bi, Fig.S1a-b**). While some limited signal was potentially discernible between PCW6-8 (**Fig.1cii**), this was not specific to the PM (**Fig.S1b**), and no quantitatively significant differences in expression within the PM were detected between any timepoints examined, including around periods of *CYP26A1* upregulation (PCW8/PCW17) (**Fig.1bi,ci**, **Fig.S1a,c**).

**Fig. 3.**
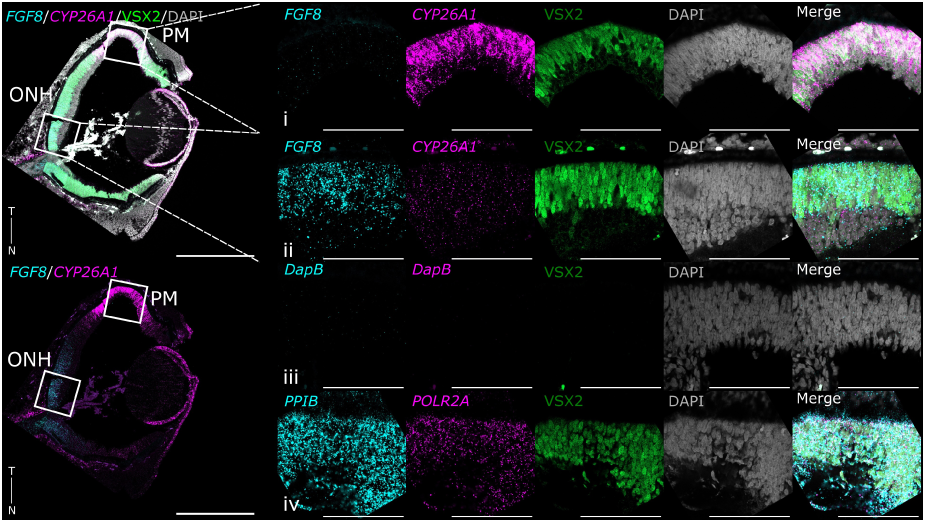
*CYP26A1* is highly expressed in the PM of early PCW8 (CS20) human fetal retina, while *FGF8* is localized exclusively to the optic nerve head (ONH). Early PCW8 (CS20) fetal retina stained *in situ* for *CYP26A1* (magenta) and *FGF8* (cyan), showing strong expression in the PM of *CYP26A1* colocalizing with RPC marker VSX2 (green), but no *FGF8*, which is visible in RPCs around the ONH. Assay negative control (*DapB*) and positive control (*PPIB* cyan, *POLR2A* magenta). Sections counterstained with DAPI (greyscale). T: Temporal; N: Nasal; CS: Carnegie Stage; PCW: Post-conception Weeks; PM: Presumptive Macula; ONH: Optic Nerve Head. Images taken at 20x (b)/40x (a). Images representative of n=3 samples. White box indicates magnified region of interest. Scale bars: (a) 500μm, (b) 100μm.

### The PM is located more peripherally in early development, relative to its central position in the mature retina, due to centre-to-periphery proliferation gradient

In the adult eye, the macula is located at the center of the retina, approximately 3.5-5.5mm temporal to the ONH(Bringmann *et al*., 2018). At early developmental timepoints, however, the spot of *CYP26A1* expression identified as the PM was more peripheral than expected (**Fig.1a**), prompting us to examine growth and proliferation across regions of the retina throughout early development.

We measured the distance between the center of the region of *CYP26A1* expression (the PM) and i) the peripheral edge of the retina; ii) the ONH (**Fig.4ai-iii**). The distance between PM-periphery increased from 300µm to 9000µm between PCW7-17, growing in a linear manner (R^2^=0.97; **Fig. 1a**, **Fig.4aiv**). The distance from PM-ONH also grew in a linear manner, from 400µm to 6000µm (R^2^=0.96; **Fig. 4aiv**), but the rate of increase was ∼2-fold lower (890µm/week and 490µm/week, respectively, *p*>0.001). Notably, while the distance PM-periphery increases as a proportion of total retina length, from 20% at PCW7 to 45% at PCW17, PM-ONH remains constant at 30% (**Fig.4av**).

**Fig. 4.**
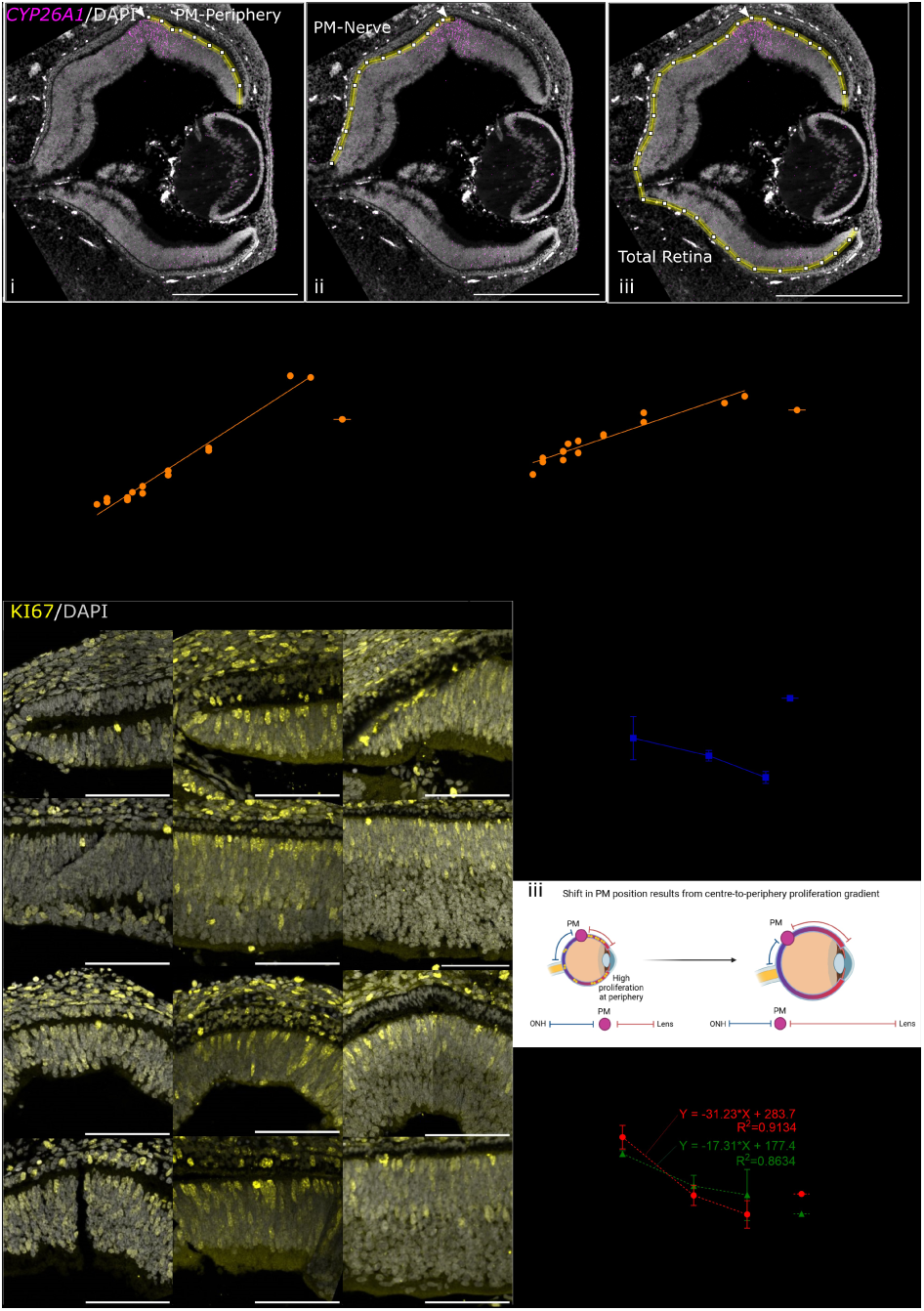
Higher proliferation in peripheral compared to central retina results in a temporally shifted *CYP26A1*+ PM at early developmental stages (PCW7-10). (a), (i-iii) The distance of the *CYP26A1*+ (magenta) PM (white arrow) to the peripheral edge of the retina/ONH. (iv) PM-periphery distance increases at a significantly higher rate (>2-fold) between PCW7-17 compared with PM-ONH distance, *p*<0.001. (v) The PM-periphery distance as a proportion of the whole retina increases from 20%-45% over PCW7-17, while PM-ONH distance remains almost constant at 30% of the whole retina (n=14, linear regression with 95% confidence internals plotted). (b) (i) Representative images of proliferative marker KI67 (yellow) staining in PCW7-8 (CS18-23) human fetal retinas shows: (ii) the proportion of DAPI+ (grey) cells expressing KI67 was at ∼two-fold higher in the peripheral compared with central retina during PCW7-8 (PCW7: *t*(2)=3.3, *p*=0.08; early PCW8 (CS20): *t*(2)=6.4, *p*=0.02; late PCW8 (CS23): *t*(2)=2.5, *p*=0.13). (iii) Schematic showing centre-to-periphery proliferation gradient, and the peripheral location of the PM in early development gradually moving towards the centre of the retina due to higher relative growth at the peripheral edge. (iv) Linear regression shows the decline in KI67+/DAPI+ cells in the PM region is significantly steeper than the nasal region between PCW7 and early PCW8, at which point *CYP26A1* is upregulated in the PM (PM: -32.2, nasal: -17.3, *p*=0.046). Region masking and full linear regression shown in **Fig.S2**. Significance values determined by paired/unpaired t-tests: \**p* value<0.05, \*\**p* value<0.01. n=3 samples/timepoint, data shown as mean±SD. PCW: Post- conception Weeks; CS: Carnegie Stage; PM: Presumptive Macula, ONH: Optic Nerve Head. Images taken at 20x. Scale bars: (a) 500μm, (b) 100μm.

To determine whether this relative positional shift of the PM was the result of variable proliferation across the retina, we examined regional proliferation using proliferative marker KI67 (**Fig.S2a**). Consistent with the known center-to-periphery developmental gradient, and comparatively higher proliferation in the periphery at later stages of retinal development(Hendrickson, 2016; Hoshino *et al*., 2017), the proportion of Ki67+ cells was ∼2-fold higher in the periphery compared to the central retina at both early and late PCW8 (early PCW8: Peripheral 68%±11.6 vs Central 36%±3.4, late PCW8: Peripheral 45%±19.1 vs Central 22%±3.7, respectively **Fig. 4bi-ii**). The central-to-periphery proliferation gradient thereby explains the more peripheral location of the PM at early stages, compared to the adult eye (**Fig. 4biii**).

### Reduction in proliferation marker KI67 in the PM coincides with *CYP26A1* expression at PCW8

We next examined proliferation within the PM during the first wave of *CYP26A1* expression, compared to a region in the nasal retina equidistant from the periphery i.e. at the same position along on the central-peripheral developmental gradient (**Fig.S2a**). As expected, both regions showed significant reductions in the proportion of proliferating cells between PCW7 and early PCW8 (PM: *t*(4)=6.5, *p*=0.003; nasal: *t*(4)=5.0, *p*=0.007; **Fig. 4biv, Fig.S2b**). However, linear regression analysis showed that the rate of reduction in the PM region was nearly twice that of the equivalent nasal region (- 31.2%/week and -17.3%/week, respectively, *p*=0.046) (**Fig.4biv, Fig.S2c**). This indicates that the reduction in proliferation in the PM goes beyond that of the central-to-peripheral gradient; moreover, this rapid reduction in proliferation in the PM coincides with the first wave of *CYP26A1* expression.

### hROs express components of RA synthesis and FGF8 pathways during early differentiation, demonstrating similarity to peripheral retina development

Our established 2D/3D differentiation protocol (**Fig.S3a**) replicates the timings of human development, yielding hROs with ∼1:4 ratio of cones:rods, a proportion similar to the outer edges of the macula(Gonzalez-Cordero *et al*., 2017). Note that this protocol includes the addition of exogenous RA from d70, which is important for maintaining retinal structure in the later stages of maturation(Sanjurjo-Soriano *et al*., 2022) but by which time cone photoreceptor neurogenesis has already occurred (**Fig.S3b-e**).

We examined the endogenous levels of RA and FGF8 signaling in our hROs. RA synthesizing enzymes, *ALDH1A1* and *ALDH1A3,* were both upregulated early in differentiation (d40/50), compared to undifferentiated hESCs (*q*(2)=13.3, *p*=0.02; *q*(2)=23.2, *p*=0.005; **Fig.5a,b**). CRABP2 traffics cytoplasmic RA to the nucleus and is concomitantly upregulated with *all*-trans-RA, making it a proxy for circulating RA levels(Napoli, 2016). *CRABP2* expression increased throughout differentiation, particularly beyond d70, when exogenous RA is added (*q*(2)=7.3, *p*=0.05; **Fig.5c**). In contrast, *CYP26A1* expression was lower in d40 hROs, compared to undifferentiated hESCs (*q*(2)=15.4, *p*=0.01) and remained low until the addition of exogenous RA at d70, whereupon, unsurprisingly, it increased (*q*(2)=8.0, *p*=0.04) and remained raised until d150 (**Fig.5d**).

**Fig. 5.**
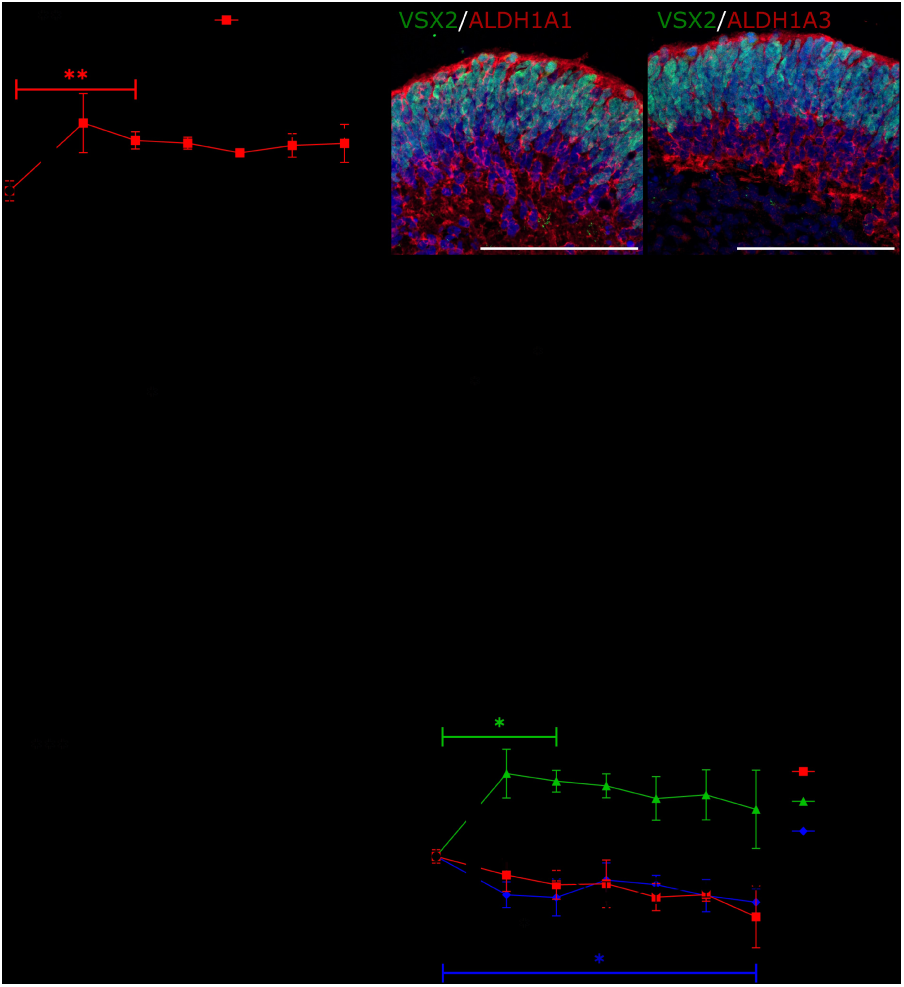
Differentiating human retinal organoids express retinoic acid and FGF8 pathway components. (a) Relative expression of RA synthesizing enzymes across hRO differentiation showing significant upregulation of *ALDH1A1* from d40 (*q*(2)=13.3, *p*=0.02), and *ALDH1A3* from d50 (*q*(2)=23.2, *p*=0.005). (b) Immunostaining of ALDH1A1 and ALDH1A3 in d40 hROs showing expression throughout the organoid, co-stained with neural retinal progenitor cell marker VSX2, counterstained with DAPI (blue). (c-f) Relative expression of mRNA for: (c) proxy for circulating RA levels *CRABP2,* with expression increasing from d40, but significantly upregulated at d80, following exogenous RA addition (*q*(2)=7.3, *p*=0.05); (d) retinoic acid catabolizing enzyme *CYP26A1*, initially downregulated compared to hESCs (*q*(2)=15.4, *p*=0.01), but upregulated at d80 following standard protocol addition of exogenous RA to culture medium from d70 (*q*(2)=8.0, *p*=0.04); (e) fibroblast growth factor 8 (*FGF8*), which has significantly increased expression at d40 (*q*(2)=18.02, *p*=0.008) and expression remains high, although slowly decreases up to d90; and (f) FGF receptors (*FGFR1*/*2*/*3*/*4*), which all showed slight downregulation compared to hESCs (*FGFR1* d60: *q*(2)=14.1, *p*=0.01; *FGFR4* d90: *q*(2)=8.2, *p*=0.04), except *FGFR3* which was significantly upregulated at d50 (*q*(2)=10.2, *p*=0.02), with expression remaining high, although slightly decreasing to d90. Further hRO marker expression shown in **Fig.S3**. qPCR data shown as Log_2_ fold change of CT values relative to d0 hESCs (n=5 pooled hROs/sample, N=3). Significance values determined by one-way repeated measures ANOVA followed by Dunnett’s test against control group, adjusting for multiple comparisons: \**p* value<0.05, \*\**p* value<0.01, \*\*\**p* value<0.001, data shown as mean±SD. Staining images representative of n=3 hROs from N=1 batch. Scale bars: 100μm.

*FGF8* was higher in d40 hROs compared to undifferentiated ESCs (*q*(2)=18.02, *p*=0.008), but expression decreased thereafter (**Fig.5e**), whilst *FGFR3* was upregulated between d40-90, (reaching significance at d50, *q*(2)=10.2, *p*=0.02), indicating FGF8 signaling is active over this period of hRO differentiation (**Fig.5f**). Expression of *FGFR1/2/4* remained low throughout (**Fig.5f**).

### Early inhibition of RA by AGN193109 leads to smaller hROs and reduced proliferation

Given the striking biphasic expression of *CYP26A1* within the human PM, we sought to mimic these periods of reduced RA signaling in hROs to better understand their potential role(s) in macula formation. In human fetal retina, the first wave of *CYP26A1* peaks in early PCW8, prior to RXRγ and RECOVERIN protein expression (**Fig.2a**), which is equivalent to d40 in hROs (**Fig.S3e**). We therefore attempted to recapitulate first wave of *CYP26A1*-induced RA catabolism by dosing hROs between d42- 50 (“early”) with pan-RAR antagonist AGN193109 (10µM) (**Fig.S4a**)(Schubert and Germain, 2023). Early dosing with AGN193109 significantly reduced expression of *CRABP2* (*t*(2)=5.77, *p*=0.03; **Fig.S5ai**) and *CYP26A1* (*t*(3)=7.91, *p*=0.004; **Fig.S5ai**), demonstrating that this dose effectively reduced levels of circulating RA. We also saw a concomitant significant increase in *FGF8* (*t*(5)=5.51, *p*=0.003; **Fig.S5aii**), indicating RA activity has been successfully inhibited, resulting in activation of downstream pathways(Cunningham and Duester, 2015).

Given the significant reduction in proliferation seen in the fetal PM during the first wave of *CYP26A1* expression (**Fig.4biv**), we examined growth of hROs dosed with RA inhibitor AGN193109 to determine whether RA inhibition reduced proliferation. Indeed, RA inhibition between d42-50 significantly reduced growth by d60, compared with DMSO-treated controls (*t*(32)=5.18, *p*<0.0001), leading to significantly smaller hROs at d150, following cessation of proliferation at 15 weeks(Gonzalez-Cordero *et al*., 2017) (*t*(16)=2.66, *p*=0.017) (**Fig.6a, Fig.S5b**). Quantification of proliferative marker KI67 also showed a modest (∼10%) but significant reduction in the proportion of KI67+ cells in AGN193109- dosed hROs at d60, compared with DMSO-treated controls (*t*(24)=2.48, *p*=0.02) (**Fig.6bi**). Conversely, there was no difference in the number of cells labelled with the apoptotic marker CASPASE-3 (*t*(29)=0.64, *p*=0.5) (**Fig.6bii**).

**Fig. 6.**
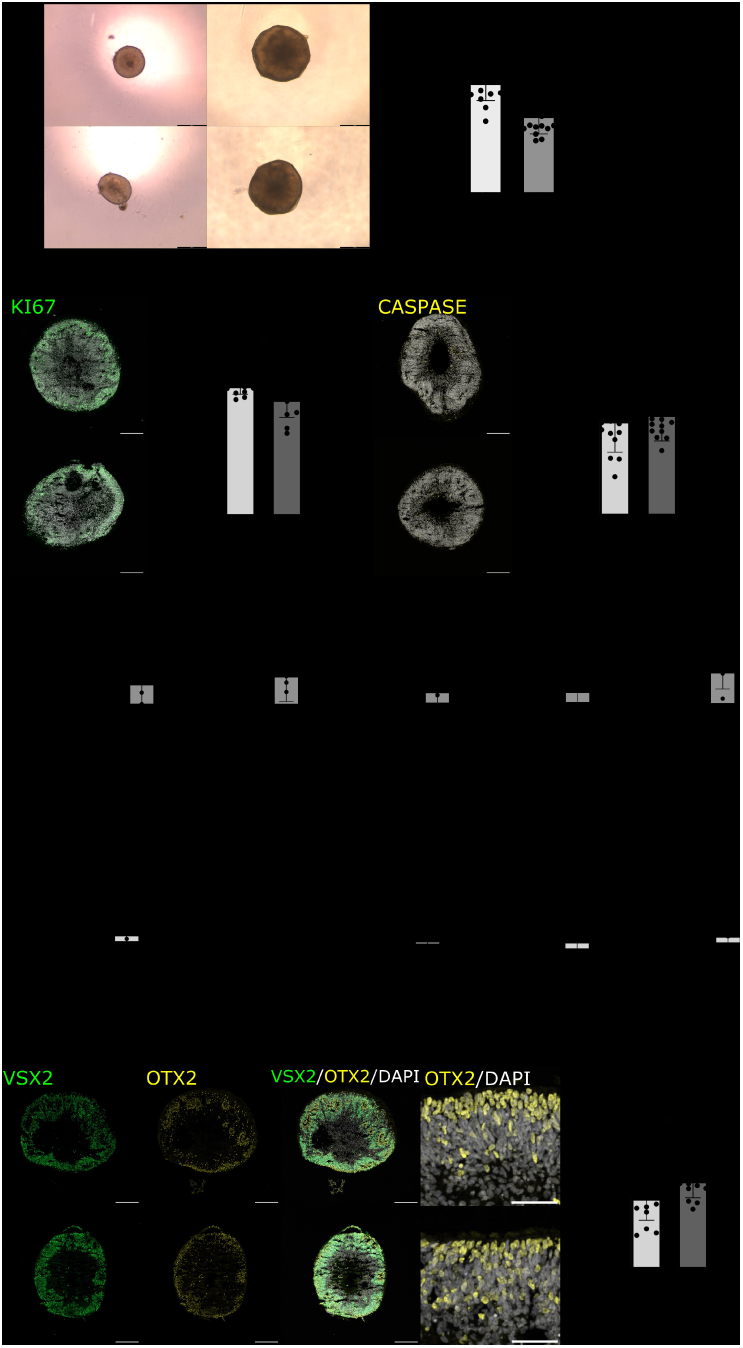
Dosing early hROs with RA receptor inhibitor AGN193109 results in reduced proliferation and increased cone precursor marker OTX2 expression. (a) Following dosing between d42-50 (“early”), the % change in area from d42-d60 was found to be significantly lower in AGN193109 dosed hROs compared with DMSO dosed controls, demonstrating reduced growth in RA inhibited hROs (t(32)=5.18, *p*<0.0001). (b), Immunostaining for (i) apoptotic marker CASPASE-3 (yellow) at d60 showed no significant increase in cell death between AGN193109 dosed and control hROs (t(29)=0.64, p=0.5), (ii) proliferative marker KI67 (green) showed a significant reduction in cell proliferation in RA inhibited hROs (t(24)=2.48, p=0.02). (c) (i) Dosing with AGN193109 led to significant increases in photoreceptor marker *RECOVERIN* (*t*(5)=2.56, *p*<0.05), and cone progenitor marker *OTX2* (*t*(4)=4.58, *p*=0.01), while early cone precursor markers *RXRγ* and *THRβ* trended towards increased expression (*t*(5)=1.45, *p*=0.2)/(*t*(5)=1.69, *p*=0.15), with no increase in retinal progenitor cell (RPC) marker *VSX2* (*t*(5)=1.39, *p*=0.2). (ii) Dosing with rFGF8 had no effect on any marker expression tested (*RECOVERIN t*(2)=0.22, *p*=0.85; *OTX2 t*(2)=2.99, *p*=0.10; *RXRγ t*(3)=0.23, *p*=0.83; *THRβ t*(2)=1.01, *p*=0.40; *VSX2 t*(3)=0.39, *p*=0.72). (d) Immunostaining of AGN193109 and DMSO dosed control hROs at d60 showed a significant increase in proportion of cone progenitor marker OTX2+ (yellow)/DAPI+ (grey) cells in AGN193109 dosed hROs (*t*(28)=2.4, *p*=0.02). Further marker expression following early dosing shown in **Fig.S5**. qRT- PCR data shown as Log2 fold change of CT values relative to DMSO treated controls (n=5 pooled hROs/sample, N≥3). Significance values determined by paired (qPCR)/unpaired (IHC) t-tests: \**p* value<0.05, \*\*\*\**p* value<0.0001, n ≥13, data shown as mean±SD. Brightfield/staining images representative of n≥13 hROs from N=2 batches. Scale bars: 200um.

### Early pharmacological inhibition of RA signaling in hROs increases expression of photoreceptor markers, while addition of recombinant FGF8 has no effect

The data above suggests that, in keeping with previous studies which show RA regulates the balance of proliferation and neurogenesis(Todd *et al*., 2018), inhibition of RA between d42-50 promotes RPC cell cycle exit in hROs. We therefore examined whether this earlier cell cycle exit increased the adoption of early born cell fates. Indeed, we saw significantly increased expression of both pan- photoreceptor marker *RECOVERIN* (*t*(5)=2.56, *p=*0.05) and cone progenitor maker *OTX2* (*t*(4)=4.58, *p*=0.01) at d60 (**Fig.6ci**). The proportion of OTX2+ cells was also ∼25% higher in early AGN193109- dosed hROs than controls (*t*(28)=2.4, *p*=0.02) (**Fig.6d**). No significant changes were observed in the expression of RPC marker *VSX2* (**Fig.6ci,d, Fig.S5di**), cone progenitor markers *RXRγ*/*THRβ* (**Fig.6ci, Fig.S5dii**) or RGC marker *RBPMS* and RGC/bipolar/amacrine marker *ISLET1* (**Fig.S5ci-ii, diii**) by d60. We also tested whether FGF8 signaling alters cone fate, as observed in chick, through addition of recombinant (r)FGF8 (100ng/ml) between d42-50 (**Fig.S4b-c**) but saw no changes in RPC or photoreceptor marker expression, compared with controls (**Fig.6cii**).

### Late RA inhibition results in decreased *S-OPSIN* and increased *M/L-OPSIN*, indicating a role in cone subtype specification

The data above indicates that the early wave of *CYP26A1* acts to promote cell cycle exit and the acquisition of a cone progenitor fate. We next sought to additionally mimic the second, later wave of *CYP26A1* expression. The “early+late” AGN193109 dosing protocol comprised of the “early” pulse at d42-50, as above, and two later pulses, the first at d70-80, when RXRY/OTX2/RECOVERIN+ hRO cone precursors have already been specified (**Fig.S3e**), equivalent to the start of the late *CYP26A1* expression wave at PCW12 (**Fig.1, Fig.2b**), and the second at d120-130, when hRO cones have matured and express *ARR3*, and rods express *RHODOPSIN* (**Fig.S3f-h**), equivalent to high *CYP26A1* expression at PCW17 (**Fig.1, Fig.2c**). Note that we used two pulses, rather than sustained application, of AGN193109 in the later phase, as continuous removal of RA between d65-120 has previously been shown to cause loss of stratification in hROs(Sanjurjo-Soriano *et al*., 2022).

No change in pan-cone marker *ARRESTIN-3* or rod marker *RHODOPSIN* gene/protein expression was observed at d150 with either early-only or early+late dosing (**Fig.7ai-ii, bi-ii**), indicating the overall proportions of cones and rods were not affected. However, those receiving early+late dosing showed a significant 2-fold reduction of *S-OPSIN* (*t*(5)=6.68, *p*=0.001) and a concurrent, significant ∼4-fold increase in *L-OPSIN* (*t*(5)=3.27, *p*=0.02) and ∼2-fold increase in *M-OPSIN* (*t*(5)=1.53, *p*=0.04), compared to DMSO controls (**Fig.7aiii-v**). Conversely, hROs receiving early-only dosing showed no changes in cone subtype marker expression at d150 (**Fig.7aiii-vi)**. Immunostaining analysis revealed that early+late dosing with AGN193109 led to a significant reduction in the proportion of ARR3+ cones expressing S-OPSIN protein (*t*(11)=3.14, *p*=0.01) (**Fig.7biii,v**), although we did not observe a change in the proportion of M/L-OPSIN/ARR3+ cones (**Fig.7biv,vi**). These results indicate that inhibition of RA signaling later in cone development alters cone subtype specification.

**Fig. 7.**
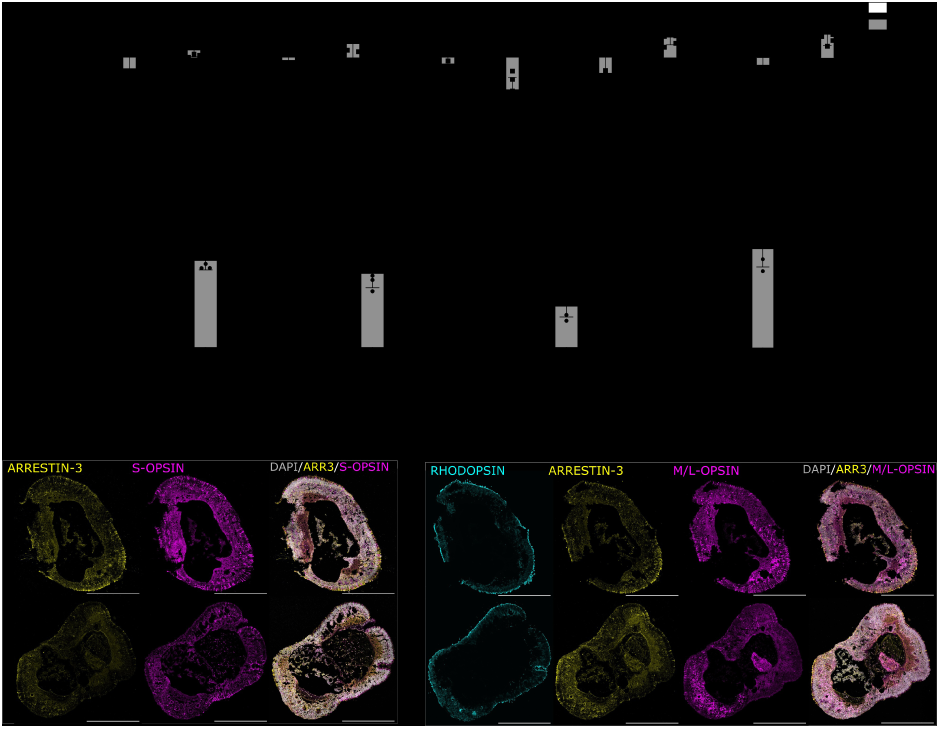
Additional late dosing of hROs with RA receptor inhibitor AGN193109 results in increased M/L- OPSIN and reduced S-OPSIN expression, indicating shift in cone subtype specification towards fovea- like composition. (a) Graphs show the effects of dosing with RA receptor inhibitor AGN193109 early in differentiation (“early”; d42-d50), or “early+late” (d42-50 and d70-80/d120-130) on photoreceptor mRNA expression at d150. Early-only dosing had no effect on any marker expression tested: (i) *ARRESTIN*-3 *t*(4)=1.80, *p*=0.15; (ii) *RHODOPSIN t*(4)=0.63, *p*=0.56; (iii) *S-OPSIN* (*t*(4)=1.06, *p*=0.35); (iv) *M*-*OPSIN t*(4)=2.3, *p*=0.08); and (v) *L-OPSIN,* (W=9, n=5, *p*=0.31). Early+late dosing lead to (i) no change in *ARRESTIN*-3, (W=-15, n=6, *p*=0.16); (ii) or *RHODOPSIN* (t(5)=2.48 *p*=0.06); (iii) a significant decrease in *S-OPSIN* (*t*(5)=6.68, *p*=0.001), (iv) a significant 1.8-fold increase in *M*-*OPSIN* (*t*(5)=2.82, *p*=0.04) and (v) a significant 3.5-fold increase in *L-OPSIN,* (*t*(5)=3.27, *p*=0.02). (**b**) Addition of late pulses of AGN193109 (early+late) led to no change in proportion of DAPI+ (grey) cells expressing (i) ARRESTIN-3 (*t*(11)=1.29, *p*=0.22) or (ii) RHODOPSIN (*t*(11)=0.41, *p*=0.69), (iii) but a significant decrease in the proportion of ARRESTIN-3+ cells expressing (iv) S-OPSIN (*t*(11)=3.14, *p*=0.01), (iv) although no change was observed in the proportion of ARRESTIN-3+ cells expressing M/L- OPSIN (*t*(11)=0.14, *p*=0.89). qRT-PCR data shown as Log_2_ fold change of CT values relative to DMSO treated controls (n=3 pooled hROs/sample, N≥5 samples/group). Significance values determined by paired (qPCR)/unpaired (IHC) t-tests or Wilcoxon test: \**p* value<0.05, \*\**p* value<0.01, data shown as mean±SD. Staining data representative of n>5 hROs from N=2 batches. Scale bars: 500μm.

## Discussion

Analysis of early-stage human macular development has been very limited to date, in particular prior to PCW8, with small sample sizes due to scarcity. The earliest previous *in situ* spatial analysis of *CYP26A1* expression in human fetal retina was late PCW8 (CS22-23), which showed expression in the PM at CS22 but none at CS23(Silva and Cepko, 2017). Here, we provide the first spatiotemporal developmental timeline of *CYP26A1* expression in the developing human macula, revealing a striking biphasic pattern of expression. Importantly, we show that the first wave of *CYP26A1* expression is initiated in the RPCs of the PM at PCW7, prior to cone genesis, and is already reducing by late PCW8, while the second wave is evident from PCW12 and is specific to the MG within the INL of the PM. This second wave in the MG layer is in line with transcriptomic studies on adult human retina(Peng *et al*., 2019; Cowan *et al*., 2020), as well as a recently published single-nuclei gene expression data of PCW8- 23 fetal retina, which showed *CYP26A1* was highly enriched in PCW17 macula cells(Zuo *et al*., 2024). Our findings are also consistent with a recent report of *CYP26A1* expression in rhesus macaque development(Krueger *et al*., 2024), which presented strong, biphasic *CYP26A1* expression in the PM region(Hendrickson and Zhang, 2019). In both human and macaque, *CYP26A1* was only expressed in RPCs during early macular formation, with no *CYP26A1* mRNA observed in the OTX2+/RXRγ+/RECOVERIN+ cone precursor cells of the ONL, indicating that RA inhibition may play a role in progenitor cell differentiation.

In contrast to HAA formation in the chick, where there is a striking spatiotemporal pattern of *Fgf8* expression in the HAA, we saw no indications of focal *FGF8* expression specific to the *CYP26A1+* PM. Instead, *FGF8* expression remained low and exhibited no significant changes in expression in the human PM throughout PCW6-17 of development. Specifically, *FGF8* was not upregulated following *CYP26A1* expression in the PM at PCW8, in contrast to the chick HAA(da Silva and Cepko, 2017), suggesting additional/different regulatory mechanisms downstream of *CYP26A1* and low RA in the human macula. These findings are consistent with recent human fetal single-nuclei gene expression data, which found *FGF8* was enriched in peripheral, but not macula, RPCs(Zuo *et al*., 2024), and RNAscope analysis of rhesus macaque development, where *FGF8* was not enriched in the developing macula region at any timepoint examined(Krueger *et al*., 2024). Moreover, we found addition of rFGF8 to hRO cultures had no impact on photoreceptor marker expression. These suggest that *FGF8* is unlikely to play a significant role in human macula formation or macula cone differentiation. They also indicate a degree of divergence in the molecular mechanisms underlying high-acuity region development in different species and particularly warrant further investigation into the pathways downstream of *CYP26A1* in the human retina.

RPC differentiation and maturation occurs earlier in the macula than the rest of the retina, shown previously by primate and human studies(Vail, Rapaport and Rakic, 1991; Hendrickson, 1992; Provis and Hendrickson, 2008; Lu *et al*., 2020; Sridhar *et al*., 2020). Whilst peripheral RPCs have been shown to maintain their progenitor state for longer than those in the PM, and higher rates of proliferation are observed in the periphery from PCW8(Hendrickson, 2016; Hoshino *et al*., 2017), earlier region-specific differences have not been studied due to lack of markers and access to tissue. Previously, bulk retina RNAseq showed enrichment of proliferation-related genes between PCW7-10, and immunohistochemistry showed reduced KI67 expression in the central compared to peripheral retina at PCW8(Hoshino *et al*., 2017). Developing this further, we found that there is a marked reduction in proliferation specifically within the PM at early PCW8 (CS20-21), coincident with the first wave of *CYP26A1* expression. Given the known role of RA in maintaining the cell cycle(Kam *et al*., 2012), this first wave of *CYP26A1* appears to serve to promote early cell cycle exit within the PM.

We investigated *CYP26A1’s* roles in human macular development by inhibiting RA signaling in hROs via pharmacological blocking of RARs to mimic the low RA environment of the *CYP26A1+* PM. Given the coincident expression of *CYP26A1* and reduced proliferation in PCW7-8 PM, we asked whether early inhibition of RA encourages cell cycle exit and neurogenesis. Indeed, we observed significantly fewer proliferating cells and reduced growth in hROs exposed to early RAR inhibition, leading to significantly smaller hROs at both d60 and d150. This aligns with our previous data investigating the effects of exogenous RA on mouse ESC-derived retinal organoids (mROs), where the inverse occurs, and a short pulse of RA from d14-16 (cone precursor stage) led to an increase in proliferation and overall cell yield(Kruczek *et al*., 2017). In the current study, early-dosed hROs also displayed a significant increase in the number of OTX2+ cells at d60, consistent with early cell cycle exit leading to differentiation of early-born cell types in the forming macula. However, this increase at d60 was not reflected in a concomitant increase at d150, presumably because supplementary pathways not active in the culture system, such as ONECUT1-mediated induction of thyroid signaling factors(Emerson *et al*., 2013; Eldred *et al*., 2018), are required for complete maturation into adult cones in hROs. Similarly, while the upregulation of *FGF8* in response to early RA-inhibition in hROs (Fig.S5aiii) is consistent with RA’s known role in inhibiting *Fgf8* expression(Cunningham and Duester, 2015), it implies that in the human macula itself additional regulatory mechanisms may act to prevent *FGF8* upregulation in response to low RA concentration.

Additional pulses of RA inhibition, aimed at mimicking the second wave of *CYP26A1* expression, did not lead to a change in overall proportions of cones:rods in mature hROs, despite RA being known to promote rod production(Osakada *et al*., 2008), and removal of RA leading to more cone-rich organoids(Sanjurjo-Soriano *et al*., 2022). This may be because a more sustained period of low RA is required for a shift towards cone-rich hRO, since *in vivo CYP26A1* expression increases from PCW12 onwards and remains elevated for many weeks. As noted earlier, however, we did not try to replicate this since continuous removal of RA between d65-120 has previously been shown to cause loss of stratification in hROs(Sanjurjo-Soriano *et al*., 2022).

Cone subtypes are determined by two fate check points: first, between S or M/L fate, and if the latter, between M- or L-OPSIN expression. The foveola at the center of the macula contains only M/L-cones while S-cones are present in the periphery of the retina. S-OPSIN-expressing cones are initially present in the early PM, and it is hypothesized that they later convert to M/L-cones, indicating plasticity of cone fate(Xiao and Hendrickson, 2000; Cornish *et al*., 2004). When RA signaling was inhibited early in differentiation, prior to cone neurogenesis, we observed no change in cone subtypes at d150. However, the addition of a later period of RA signaling inhibition led to a significant upregulation of *M/L-OPSIN* and concomitant downregulation of *S-OPSIN* gene expression and fewer S-OPSIN+ cells. This reflects a recent BioRxiv report showing that *CYP26A1*-null hROs favor S-cone formation(Hussey *et al*., 2023). Interestingly, a similar effect was seen in *VAX2* knockout mice; normally *VAX2* restricts the expression of *CYP26A1* to a stripe in the central retina. In its absence, there is an expansion of M- cones into the usually S-cone dominated ventral retina(Alfano *et al*., 2011). Moreover, addition of exogenous RA to mROs after the period of cone precursor specification led to upregulation of S-opsin but no change in Arrestin-3 expression, indicating that alterations in RA signaling later in development also influence S- vs M/L-opsin cone subtype specification in mouse development(Kruczek *et al*., 2017).

RA has also recently been shown to regulate the M- vs L-cone cell fate(Hadyniak *et al*., 2024). In our hands, while the addition of late pulses of RA inhibition in hROs led to an increase in both *M-* and *L- OPSIN* expression, we saw a greater impact on *L-OPSIN* expression, compared to *M-OPSIN* (∼4-fold, versus ∼2-fold, increase respectively), indicating that lack of RA signaling more strongly favors L-cone fate. This broadly aligns with a recent report by Johnston and colleagues indicating that sustained application of RA to hROs during the same period of differentiation (d43-d130) of RA suppresses L- cone formation, whilst promoting M-cone fate(Hadyniak *et al*., 2024).

Together, these data show for the first time the spatiotemporal biphasic pattern of expression of *CYP26A1* within the PM across human macula development. Conversely, *FGF8* expression shows no such spatiotemporal changes in the PM, consistent with the macaque but in contrast to the chick HAA, suggesting a divergence of regulatory mechanisms in high-acuity area formation. Based on our analysis of RA pathway manipulation in hRO models, we propose different roles for the two waves of CYP26A1- mediated RA inhibition in macular formation: the early wave (PCW7-8) stimulates RPCs to exit the cell cycle, encouraging cone generation, while the second, later wave (from PCW12) plays a role in the specification of cone subtypes.

This work bridges a gap in fundamental knowledge of the molecular mechanisms underlying human macular development and highlights the importance (and challenges) of studying human development in both animal models and human-derived tissue. These results open new avenues for future research including resolving the molecular network downstream of *CYP26A1* and determining the drivers of *CYP26A1* expression in the human PM and how they are initiated. Elucidating these molecular mechanisms could provide strategies to encourage formation of more macular-like cellular structure in *in vitro* models to study macular development and disease, provide a preclinical model for drug testing to treat macular degeneration, and improve the efficacy of generating cells used for transplantation therapies that to restore visual function in patients.

## Experimental Procedures

### Human fetal retina sample collection

Human fetal retina samples were collected from the Human Developmental Biology Resource with research ethics approval and informed consent (Project 200599, REC 18/LO/0822 and 18/NE/0290, IRAS: 244325 and 250012). Samples were stored in compliance with the Human Tissue Act 2004. Staging was performed by the HDBR according to defined criteria (https://hdbratlas.org/staging-criteria/carnegie-staging.html). A minimum of 2 biological replicates from independent samples were analyzed for each timepoint.

### Differentiation of retinal organoids

Cells from the H9 ESC line were maintained and used to generate human retinal organoids (hROs) according to a previously published protocol(Gonzalez-Cordero *et al*., 2017). Full protocol and reagents can be found in supplementary material. For dosing, 10µM AGN193019 (Bioteche: 5758) or 100ng/µl rFGF8 (Biotechne: 423-F8-025) were added to culture media every 2 days throughout the dosing windows.

### RNAscope^TM^ *in situ* hybridization

RNAscope™ Multiplex Fluorescence Assay was performed using manufacturer’s protocol for fixed frozen tissue with manual antigen retrieval. Sections were incubated with catalogue probes Hs- CYP26A1-C1 (ACD Bio: 487741) and Hs-FGF8-C2 (ACD Bio: 415791-C2), counterstained with VSX2 and DAPI. Full protocol, reagents and quantification can be found in supplementary material.

### Immunohistochemistry (IHC)

IHC was performed using primary/secondary antibodies listed in **Table S1**/**S2**. Full protocol, reagents and quantification can be found in supplementary material.

### qRT-PCR gene expression analysis

qRT-PCR was performed to analyze gene expression using primers listed in **Table S3**. Full protocol, reagents and quantification can be found in supplementary material.

### Fetal retina regional distance measurements

Regional length measurements were taken from confocal RNAscope^TM^ images with fluorescent *CYP26A1* staining. The outer edge of the retina was manually traced using the segmented line drawing tool in ImageJ, from the center of *CYP26A1* expression (white arrow, Fig.4a) to the peripheral edge, or edge of optic stalk, and distance calculated using the measure function. Growth rates were analyzed using linear regression analysis in GraphPad Prism (v10.1.2).

### Statistics

All data are presented as mean±SD; N denotes number of independent experiments (i.e., differentiation batches) and n denotes number of images or hROs examined. Statistical testing was performed in GraphPad Prism (version 10.1.2). Tests used to analyze statistical significance are specified in methods/figure legends. Outliers were identified and excluded using the ROUT test (Q=10%). Shapiro-Wilk Tests were used to assess normality and F-tests were used to confirm equal variances prior to applying parametric statistical tests.

### Resource Availability

Further information and requests for resources/reagents should be directed to the corresponding authors.

## Materials Availability

This study did not generate new unique reagents.

## Data and Code Availability

This paper did not generate original code. Additional data is available from the corresponding authors upon request. FAIR data management principles were followed throughout.

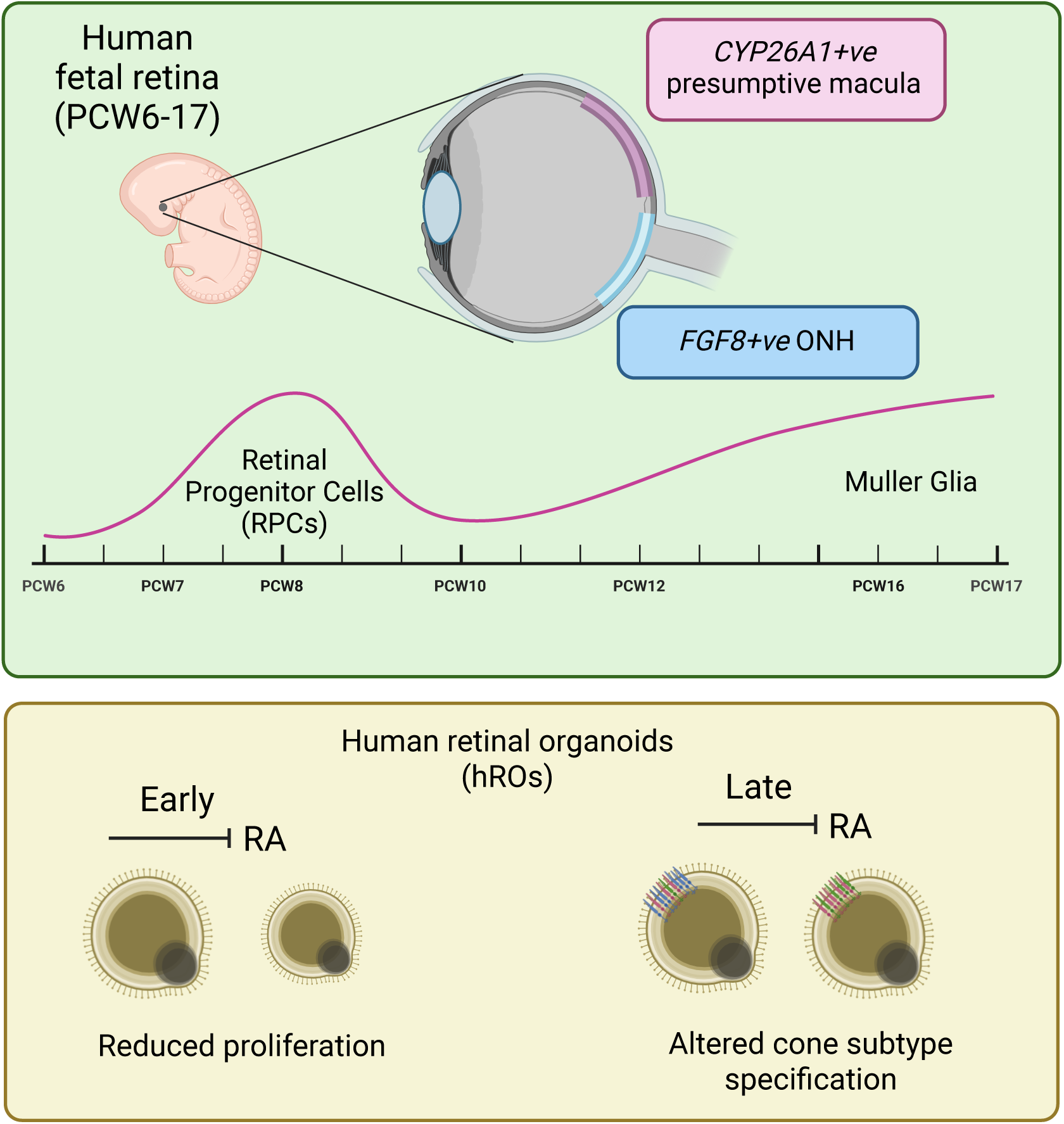

## Acknowledgements

This research was funded by Fight for Sight (5139/5140) and Medical Research Council UK (MR/T002735/2). Maja Wojtynska was funded by the Wellcome Trust as part of the Advanced Therapies for Regenerative Medicine Wellcome Trust PhD Program (218461/Z/19/Z). This work was made possible by the support, dedication and teamwork of the Ocular Cell and Gene Therapy team, with particular thanks to Dr E West, Dr M Branch, Dr M Khazim, B Ladino, J Kapadia, E Lanning, M Margari, M Tariq, C Mofidi, K Kumar, S Guilfoyle, S van Heerden for help with stem cell maintenance cultures. We wish to thank Dr M Riabiz (KCL) for providing independent expert statistical analysis advice. For the purpose of open access, the authors have applied a Creative Commons Attribution (CC BY) license to any Author Accepted Manuscript version arising. Graphical images were created in BioRender.

## Author Contributions

Conceptualization – PH, RP; Data curation – PH, MW; Formal Analysis – PH; Funding acquisition – RP; Investigation – PH, RP; Methodology – PH, RP; Supervision – RP; Visualization – PH; Writing – original draft – PH; Writing – review & editing – PH, AS, RA, RP

