## Supplementary figures and images for "Human macula formation involves two waves of retinoic acid suppression via *CYP26A1* that modulate cell cycle exit and cone subtype specification"

### Supplementary Figure 1

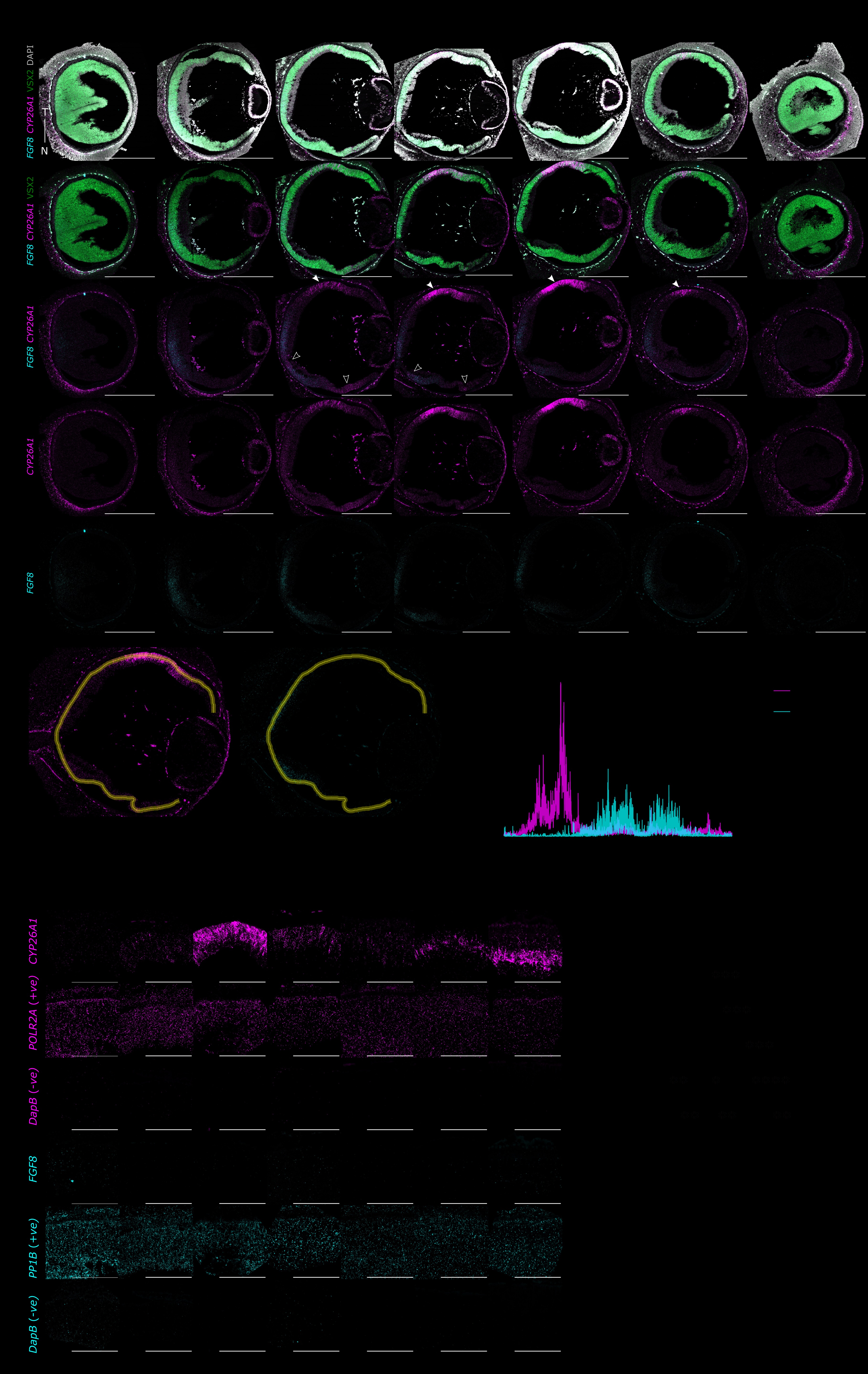

### Supplementary Figure 2

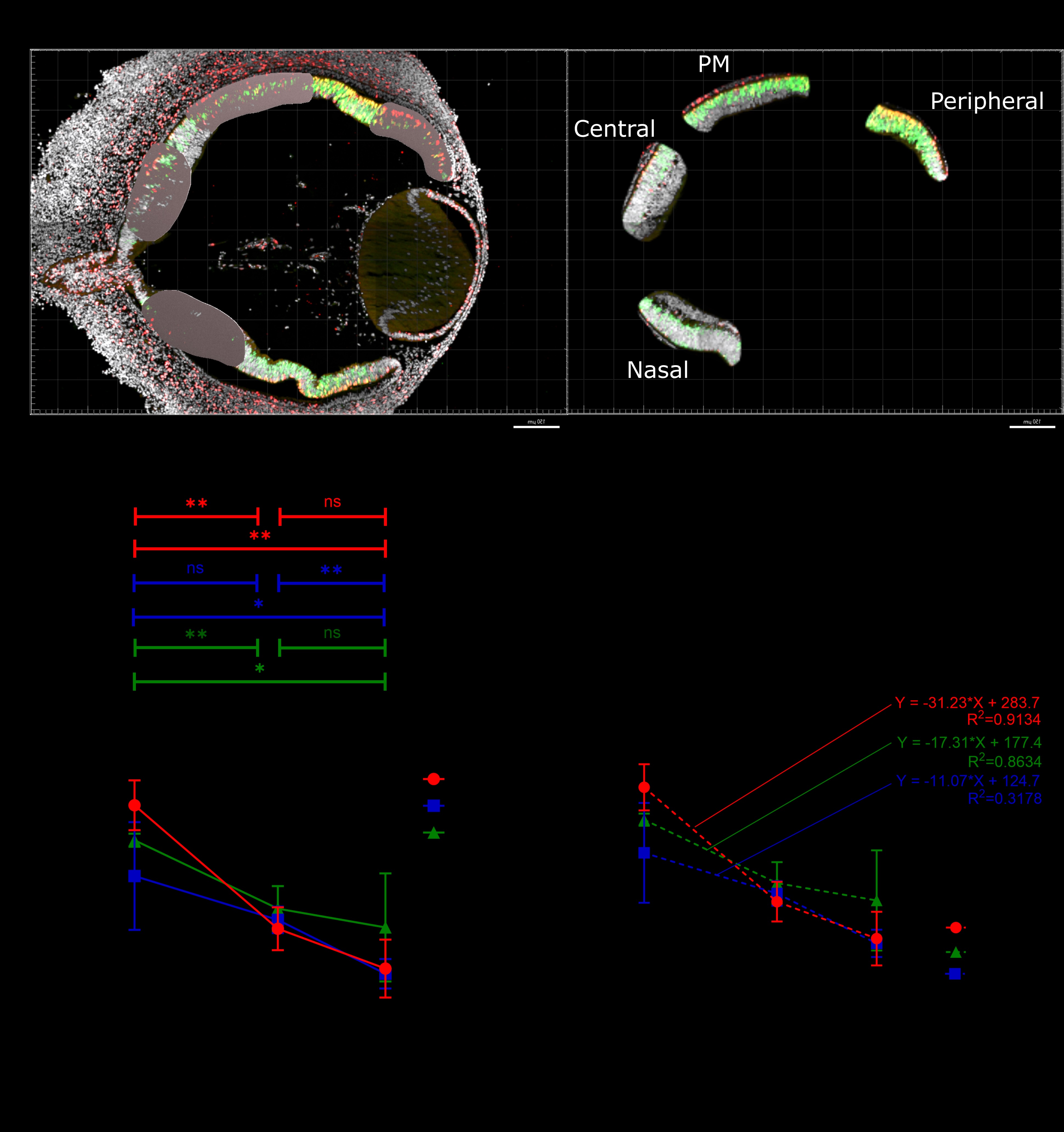

### Supplementary Figure 3

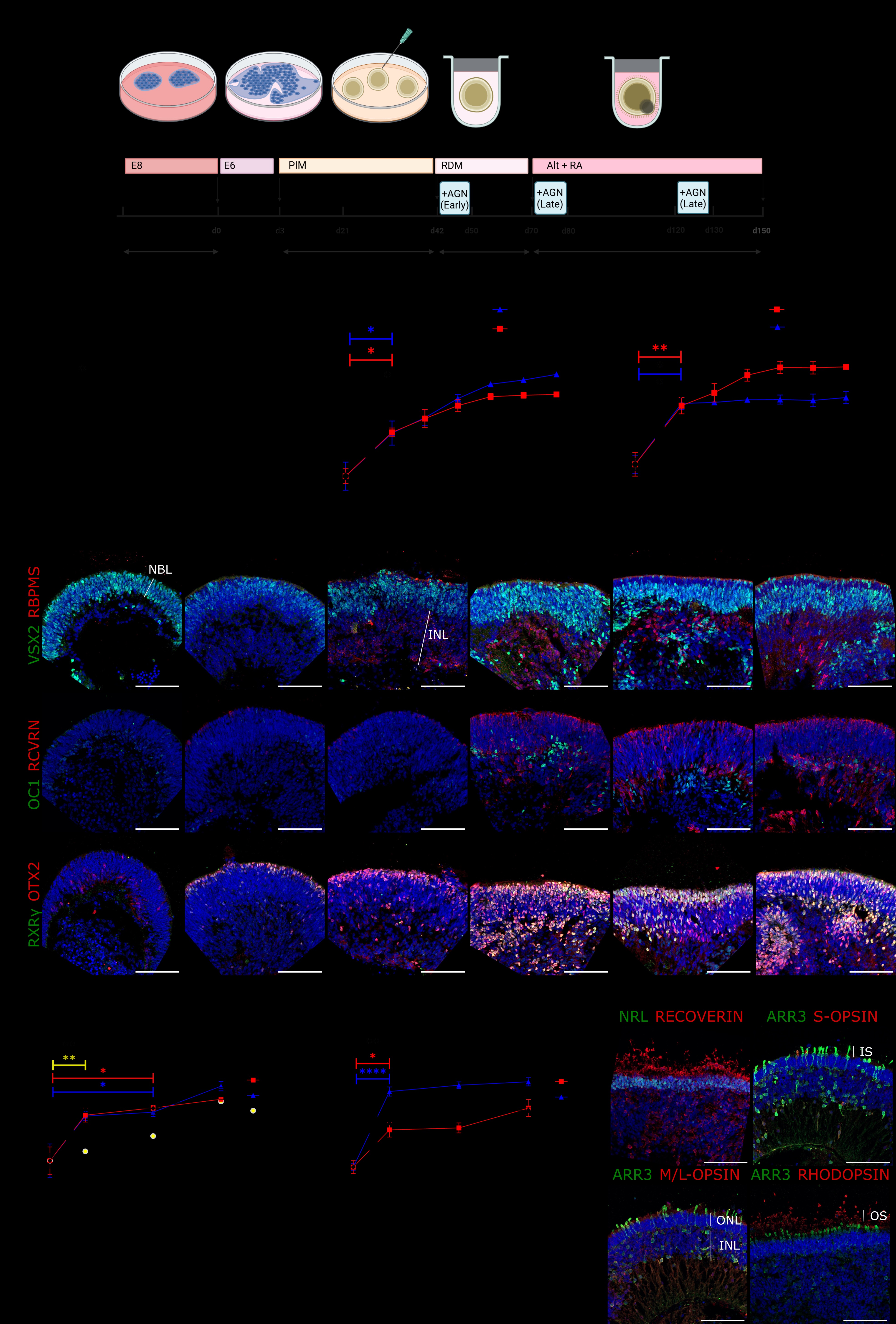

### Supplementary Figure 4

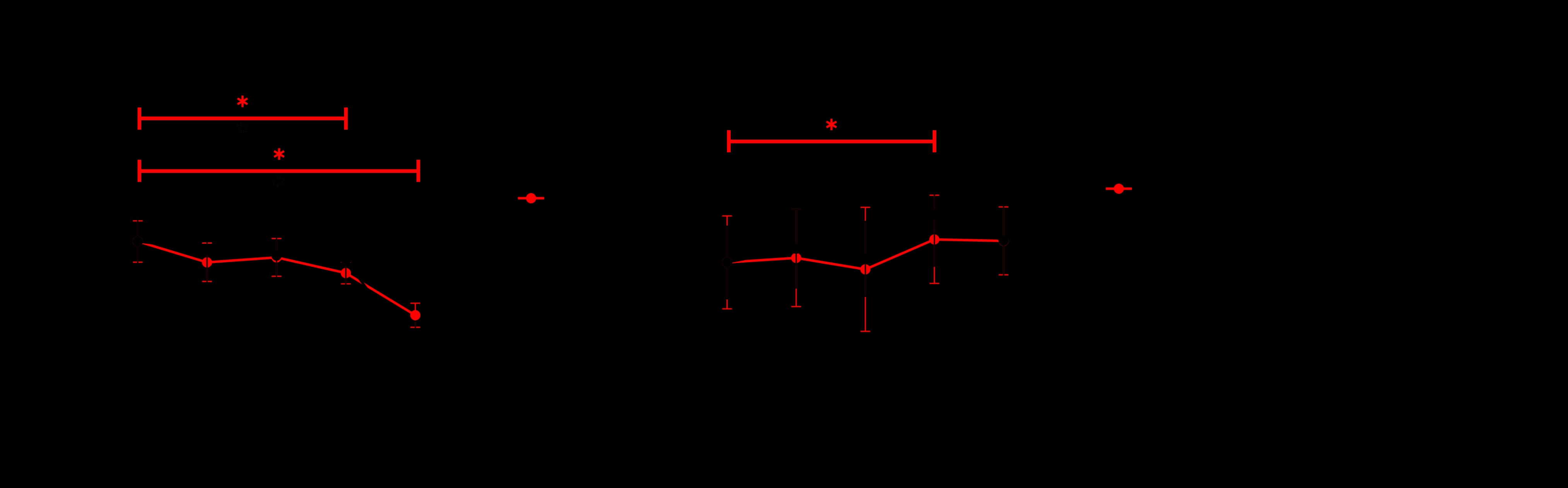

### Supplementary Figure 5

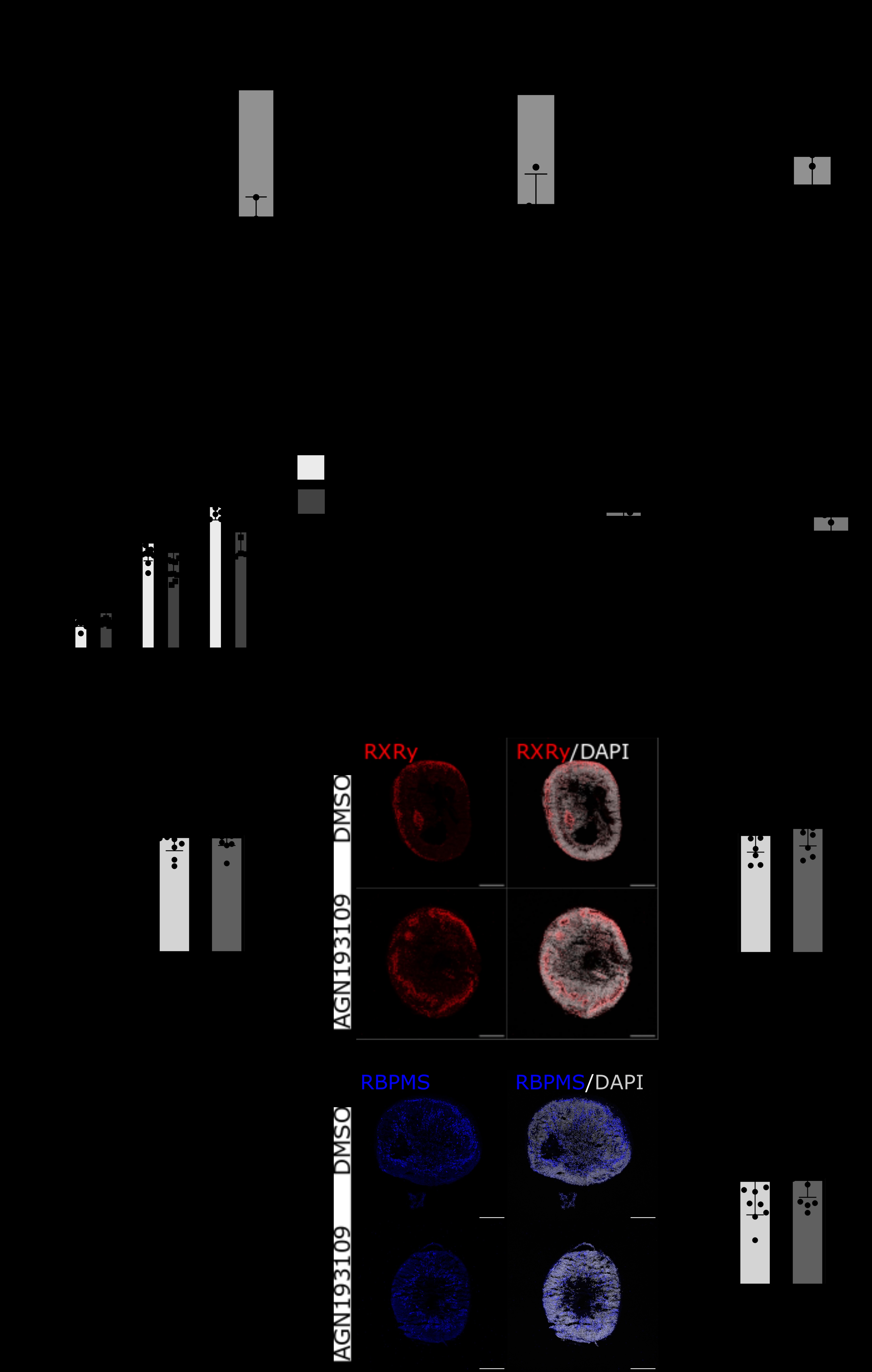

### Supplementary Figure 6

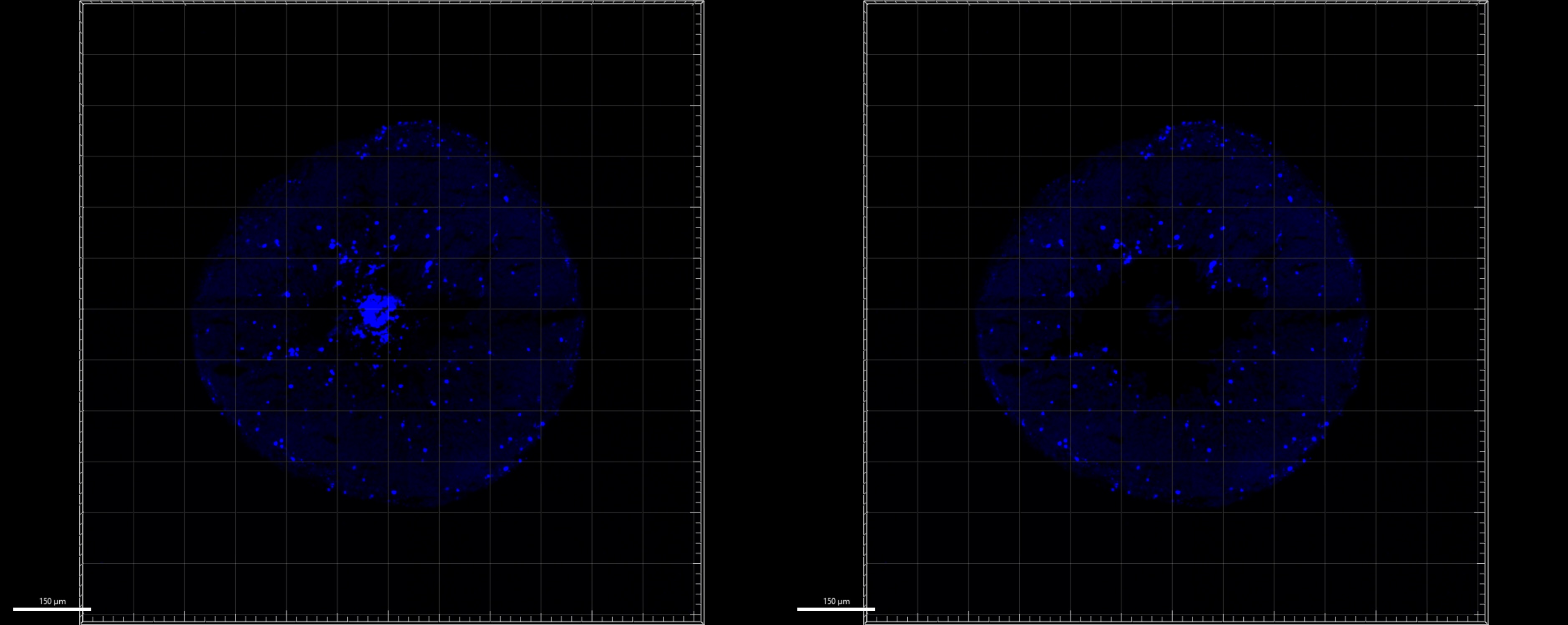
