## Supplementary Data for "Human macula formation involves two waves of retinoic acid suppression via *CYP26A1* that modulate cell cycle exit and cone subtype specification"

### Supplementary Figures

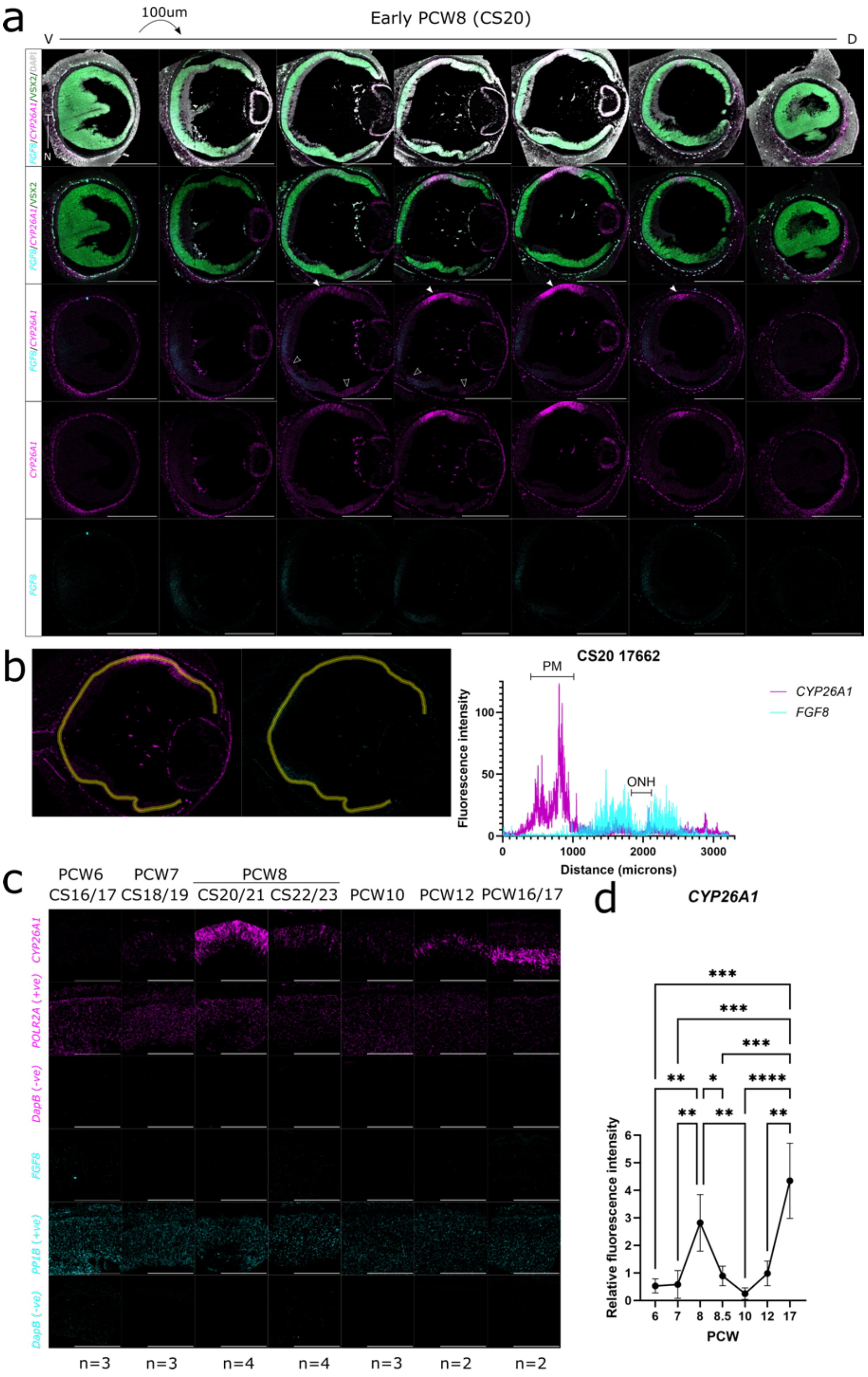

**Fig.S1. *CYP26A1* expression is localized to a single spot-like region at the presumptive macula (PM), with expression peaks at early PCW8 (CS20-21) and from PCW12 onwards, while *FGF8* is exclusively expressed around the optic nerve head, related to Fig.1.** (a) Representative images of serial sections through a CS20 human retina, taken at 100µm intervals and stained for the RA catabolizing enzyme, *CYP26A1* (magenta), and growth factor *FGF8* (cyan) mRNA using RNAscope™. Imaging shows expression of *CYP26A1* in the PM with no overlapping FGF8 expression, which is instead expressed around the optic nerve head. (b) Quantification of *CYP26A1*/*FGF8* fluorescence intensity across a single retina at CS20, showing *FGF8* is expressed in the retinal regions around optic nerve head, while *CYP26A1* is specifically expressed in the presumptive macula, where there is no distinct, overlapping *FGF8* expression. (c) Representative high-magnification images of the *CYP26A1*+ PM regions at all timepoints, including assay positive (*PPIB* cyan, *POLR2A*, magenta) and negative (*DapB*) controls showing consistent background/housekeeping expression across biological samples. (d) Quantification of *CYP26A1* normalized fluorescence intensity in the PM including all statistically significant values based on Tukey's multiple comparisons tests following one-way ANOVA. *$p$ value<0.05, **$p$ value<0.01, *** $p$ value<0.001, **** $p$ value<0.0001. Data are represented as mean ±SD. White arrows indicate PM, while white outlined arrow heads indicate low levels of *CYP26A1* expression visible around optic nerve head (ONH) and at nasal edge of the retina. Sections were co-stained with RPC marker, VSX2 (green) and counterstained with nuclear marker, DAPI (greyscale). S – Superior; I – Inferior; T – Temporal; N – Nasal; CS – Carnegie Stage; PCW – Post-conception weeks, PM – Presumptive macula. Images taken at 20x magnification. Images are representative of n=3 CS20/21 samples. Scale bars - 500µm.

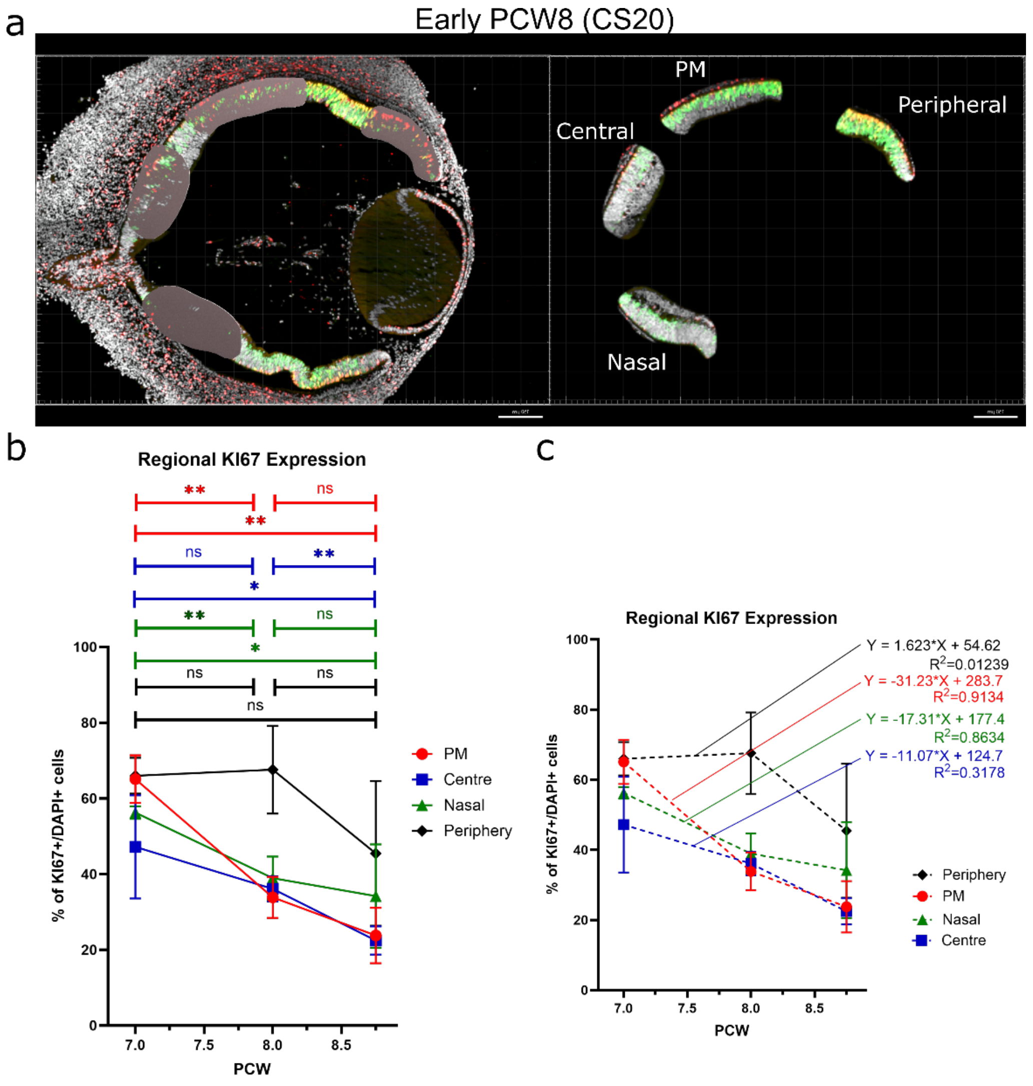
a
Early PCW8 (CS20)
PM
Peripheral
Central
Nasal
b
Regional KI67 Expression
**
ns
**
ns
**
*
**
ns
*
ns
ns
ns
% of KI67+/DAPI+ cells
PCW
PM
Centre
Nasal
Periphery
c
Regional KI67 Expression
Y = 1.623*X + 54.62
$R^2$=0.01239
Y = -31.23*X + 283.7
$R^2$=0.9134
Y = -17.31*X + 177.4
$R^2$=0.8634
Y = -11.07*X + 124.7
$R^2$=0.3178
% of KI67+/DAPI+ cells
PCW
Periphery
PM
Nasal
Centre

**Fig.S2. Proliferation in human developing retinal regions (CS18-23), related to Fig.4.** (a), Masking of PM (based on *CYP26A1*+ staining), central (adjacent to the optic nerve), nasal (equivalent distance as PM to periphery on the central-peripheral gradient on the nasal side) and peripheral regions in Imaris 10.0. (b) Proportion of KI67+/DAPI+ cells in all analyzed developing retinal regions from CS18-23. The proportion of proliferating DAPI+ cells decreases significantly in both the PM and the equivalent nasal region on the center to peripheral gradient by early PCW8 (PM: $t(4)=6.50$, $p=0.003$; nasal: $t(4)=5.03$, $p=0.007$) as well as between PCW7 and late PCW8 (PM: $t(4)=7.4$, $p=0.002$; nasal: $t(4)=2.8$, $p=0.05$). At CS18, the PM has a similar proportion of KI67+/DAPI+ cells to the peripheral retina (mean of 65/66% respectively, $t(2)=0.33$, $p=0.78$)), while at CS20/23, the PM had similar proportion of KI67+/DAPI+ cells to the central retina (CS20: means of 34/36%, $t(2)=1.15$, $p=0.37$); CS23: means of 24/22%, $t(2)=0.38$, $p=0.74$) and significant/nearing significant difference in proliferating cells compared with nasal/peripheral retina (CS20: $t(2)=5.26$, $p=0.03$/$t(2)=5.92$, $p=0.02$; CS23: $t(2)=2.47$, $p=0.13$/$t(2)=2.76$, $p=0.11$ respectively. (c) Linear regression of proportion of KI67+/DAPI+ cells in different retinal regions between CS18 and CS20 shows that the PM region has a steeper decline in KI67+ cells than any other retinal region (PM: -15.62, central: -5.5, nasal: -8.7, peripheral: 0.8, p=0.01). PM – Presumptive Macula; PCW – Post-conception Weeks. n=3/timepoint, data are represented as mean ±SD.

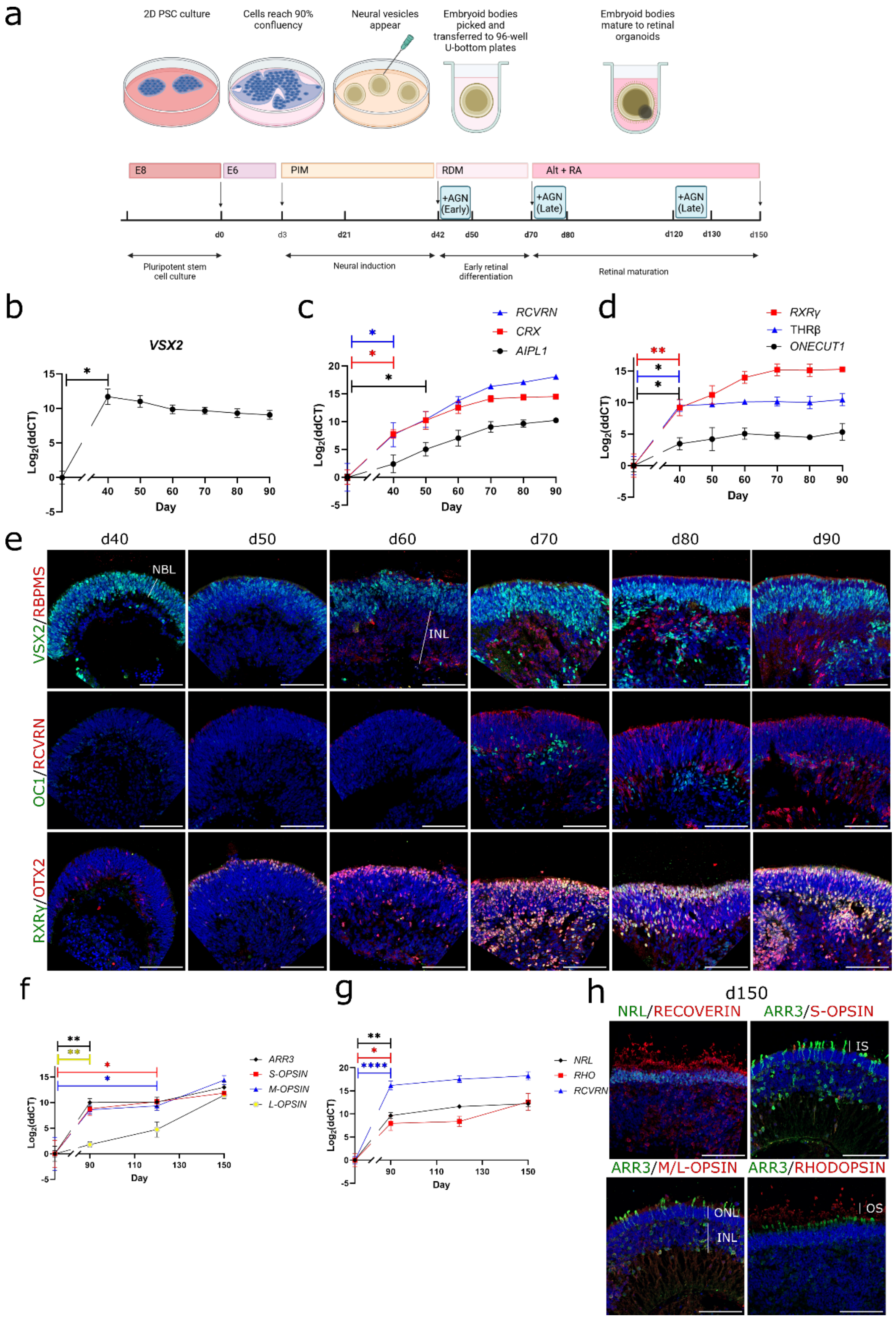

a
2D PSC culture
Cells reach 90% confluency
Neural vesicles appear
Embryoid bodies picked and transferred to 96-well U-bottom plates
Embryoid bodies mature to retinal organoids
E8
E6
PIM
RDM
Alt + RA
+AGN (Early)
+AGN (Late)
+AGN (Late)
d0
d3
d21
d42
d50
d70
d80
d120
d130
d150
Pluripotent stem cell culture
Neural induction
Early retinal differentiation
Retinal maturation
b
VSX2
Log2(ddCT)
Day
c
RCVRN
CRX
AIPL1
Log2(ddCT)
Day
d
RXRγ
THRβ
ONECUT1
Log2(ddCT)
Day
e
d40
d50
d60
d70
d80
d90
VSX2/RBPMS
OC1/RCVRN
RXRγ/OTX2
NBL
INL
f
ARR3
S-OPSIN
M-OPSIN
L-OPSIN
Log2(ddCT)
Day
g
NRL
RHO
RCVRN
Log2(ddCT)
Day
h
d150
NRL/RECOVERIN
ARR3/S-OPSIN
ARR3/M/L-OPSIN
ARR3/RHODOPSIN
IS
ONL
INL
OS

**Fig.S3. Temporal and spatial photoreceptor development in human retinal organoids, related to Fig.5.** (a) graphical representation of hRO differentiation and "early" dosing with RA receptor inhibitor AGN193019 (10µM) and recombinant FGF8 (100ng/ml) between d42-50, followed by "mid" AGN193109 pulse at d70-80 and "late" pulse at d120-130; (b-d) Relative mRNA expression of (b) RPC cell marker *VSX2* showing upregulation at d40 of differentiation compared with d0, (c) photoreceptor precursor markers, *RECOVERIN*, *CRX* and *AIPL1* showing photoreceptor specification from d40 of differentiation with expression further increasing to d90 and (d) cone precursor markers *RXRγ*, *THRβ* and *ONECUT1* significantly upregulated by d40, with plateauing expression from d60-70, indicating specification of cone precursors by this timepoint; (e) immunostaining of VSX2+ retinal progenitor cells in the neuroblastic layer (NBL) from d40 of differentiation, along with retinal ganglion cell (RGC) marker RBPMS in the inner nuclear layer (INL) from d60, OTX2+ cone-biased progenitor cells showing increasing cell numbers from d50, and RECOVERIN+ photoreceptor precursors from d70 predominantly localizing to the outer layers of developing hROs, and ONECUT1+ horizontal cell-biased retinal progenitors towards the inner layers of the organoid, alongside some OTX2+/RECOVERIN+ bipolar cells/mis-localized photoreceptor precursor cells; (f) Relative mRNA expression of mature cone markers *ARR3*, *OPSNSW*, *OPSNMW* and *OPSNLW* and (g) mature rod markers *RHODOPSIN*, *NRL* and photoreceptor marker *RECOVERIN* showing maturation of rods and cones by d150 of differentiation; (h) immunostaining of mature d150 retinal hROs stained for pan-photoreceptor marker RECOVERIN, with tight multilayered NRL+ rod nuclei in the outer nuclear layer (ONL), RHODOPSIN labelling the outer segments of rod photoreceptor cells and ARR3 labelling the cell bodies of a single apical layer of cone photoreceptor cells showing the inner segments, with a small number of S-OPSIN+ short wavelength cones, and M/L-OPSIN+ medium/long wavelength cones, with some RECOVERIN+ photoreceptors displaced to the INL. All qRT-PCR data shown as $Log_2$ fold change of CT values relative to d0 hESCs (n=3-5 pooled hROs per sample, N=3). Significance values determined by one-way repeated measures ANOVA followed by Dunnett's test against control group, adjusting for multiple comparisons: *$p$ value<0.05, **$p$ value<0.01, ****$p$ value<0.0001, data are represented as mean ±SD. PSC – Pluripotent stem cell; E8 – Essential 8 media; E6 – Essential 6 media; PIM – Proneural induction media; RDM – Retinal differentiation media; ALT – Alternative RDM; RA – Retinoic Acid; AGN – AGN193109; NBL – Neuroblastic Layer; INL – Inner Nuclear Layer; ONL – Outer Nuclear Layer; IS – Inner segments; OS – Outer segments. Staining data representative of n=3 hROs from N=2 batches of differentiation, counterstained with DAPI (greyscale). Scale bars - 100μm.

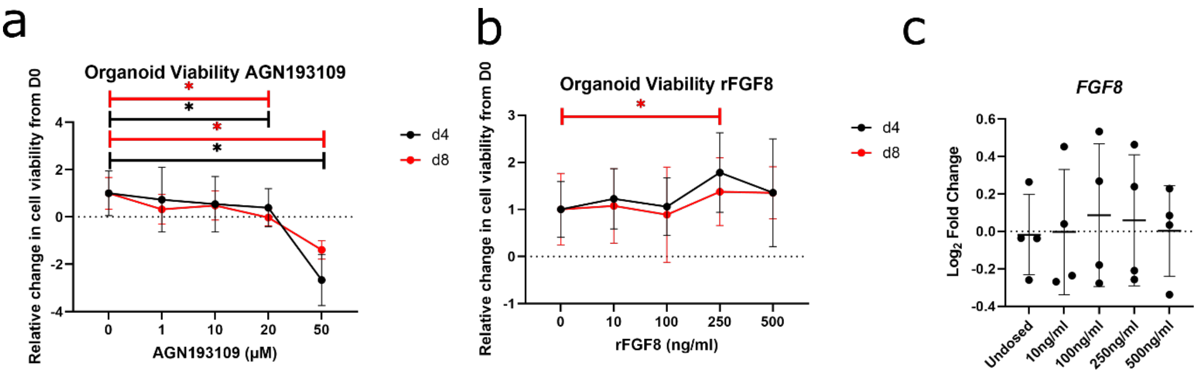


**Fig.S4. Dose optimization of AGN193109 and recombinant FGF8 for hROs based on alamarBlue cell viability assay.** Fluorescence was measured at 590nm (RFU) and change in reduction at d4 and d8, compared with d0 of dosing for each hRO, then relative change of treated compared with DMSO treated was calculated and plotted for (a) AGN193109 and (b) recombinant FGF8 (rFGF8) (n=5 from N=1 batch). (c) Dosing with rFGF8between d40-d50 lead to no significant change in *FGF8*, which is known to regulate its own expression Data shown as $Log_2$ fold change of CT values relative to PBS treated controls (5 pooled hROs per sample, N=4). Error bars - standard deviation. Significance values determined by one-way repeated measures ANOVA followed by Dunnett's test against control group, adjusting for multiple comparisons. *$p$ value<0.05.

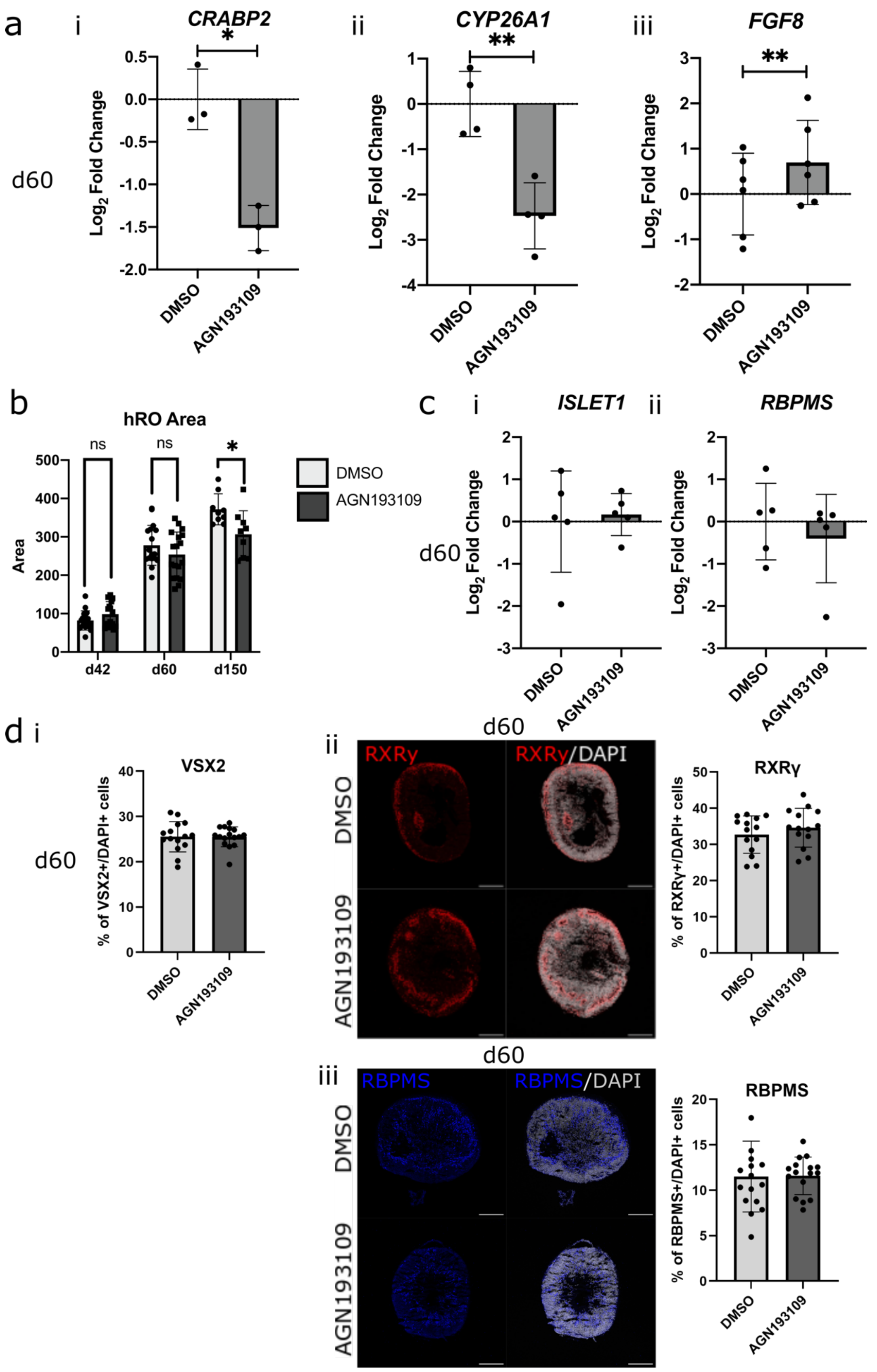

a
i
CRABP2
ii
CYP26A1
iii
FGF8
d60
Log2 Fold Change
DMSO
AGN193109
b
hRO Area
Area
ns
d42
d60
d150
c
ISLET1
RBPMS
d
VSX2
% of VSX2+/DAPI+ cells
RXRγ
RXRγ/DAPI
% of RXRγ+/DAPI+ cells
RBPMS/DAPI
% of RBPMS+/DAPI+ cells


83

**Fig.S5. Dosing early hROs with RA receptor inhibitor AGN193109 does not alter retinal ganglion cell (RGC) marker or cone precursor marker expression at d60, related to Fig.6.** (a-b) Dosing with RA receptor inhibitor AGN193109 between d42-d50 leads to (a) (i) a significant reduction in *CRABP2*, a proxy for circulating RA levels, ($t(2)=5.77$, $p=0.03$), (ii) a significant reduction in *CYP26A1,* indicating reduced cytoplasmic RA being catabolized ($t(3)=7.91$, $p=0.004$), and (iii) a significant increase in *FGF8*, a downstream effector of RA inhibition ($t(5)=5.51$, $p=0.003$); (b) Measurements of hRO area show no significant difference in size of treated/control samples at d42 (prior to dosing) (W=117, n=17, $p=0.24$) or d60 (10 days after dosing) (t(33)=1.28, p=0.21), although a significant difference in hRO growth was detected at d60 (Fig.6a), and a significant difference in hRO area was seen at d150 (t(16)=2.66, p=0.017). (c) No significant change in RGC marker *RBPMS* ($t(4)=1.02$, $p=0.37$) or RGC/bipolar/amacrine cell marker *ISLET1* (W=9, n=5, $p=0.31$). All data shown as $Log_2$ fold change of CT values relative to the mean of DMSO treated controls (n=5 pooled hROs per sample, N≥3, lines indicate paired samples of DMSO/AGN-dosed samples from the same batch of differentiation to control for batch-batch variation). Significance values determined by paired t-tests. (c) Immunostaining of AGN193109 dosed and DMSO dosed control hROs at d60 showing no significant change in the proportion of (i) RGC marker RBPMS (blue) cells ($t(30)=0.08$, $p=0.94$) (ii) cone precursor marker RXRγ (red) ($t(26)=0.96$, $p=0.35$) or (iii) RPC marker VSX2 (staining in Fig.6) ($t(29)=0.01$, $p=0.99$). Significance values determined by unpaired t-tests: *$p$ value<0.05, n>13, data are represented as mean ±SD. Staining data representative of n>13 hROs from N=2 batches of differentiation, counterstained with DAPI (greyscale). Scale bars - 100μm.

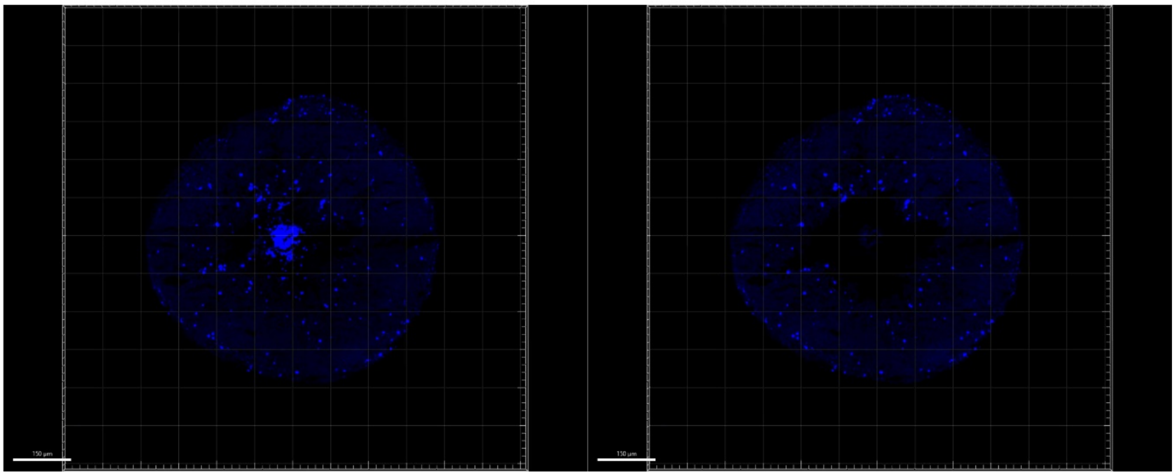

**Fig.S6. Masking of necrotic core for CASPASE-3 staining quantification in IMARIS 10.0, related to Fig.6.**

### Supplementary Tables

**Table S1 – Primary antibodies for IHC, relating to Fig.1-7, S1-3 and S5-6.**

| Target | Species | Manufacturer | Product code | RRID | Dilution |
|---|---|---|---|---|---|
| ALDH1A1 | Rabbit | Abcam | ab23375 | AB_2224009 | 1:200 |
| ALDH1A3 | Rabbit | Abcam | ab129815 | AB_2937054 | 1:200 |
| ARRESTIN-3 | Goat | Novus | NBP1-37003 | AB_2060085 | 1:100 |
| CASPASE-3 | Rabbit | Abcam | ab2302 | AB_302962 | 1:100 |
| CRX | Mouse | Abnova | H00001406-M02 | AB_606098 | 1:800 |
| KI67 | Rabbit | Abcam | ab15580 | AB_443209 | 1:100 |
| M/L-OPSIN | Rabbit | Millipore | AB5405 | AB_177456 | 1:100 |
| NRL | Goat | R+D systems | AF2945 | AB_2155098 | 1:100 |
| ONECUT1 | Mouse | Santa cruz | sc-13050 | AB_2251852 | 1:100 |
| OTX2 | Goat | R&D Systems | AF1979 | AB_2157172 | 1:50 |
| RBPMS | Rabbit | PhosphoSolutions | 1830 | AB_2492225 | 1:150 |
| RECOVERN | Rabbit | Chemicon | AB5585 | AB_2253622 | 1:1000 |
| RHODOPSIN | Mouse | Sigma | O4886 | AB_260838 | 1:1000 |
| RXRγ | Mouse | Santa Cruz | sc-365252 | AB_10850062 | 1:200 |
| S-OPSIN | Rabbit | Millipore | AB5407 | AB_177457 | 1:200 |
| SOX9 | Rabbit | Sigma | AB5535 | AB_2239761 | 1:200 |
| VIM | Chicken | Novus | NB300 | AB_922758 | 1:600 |
| VSX2 | Mouse | Santa Cruz | sc-365519 | AB_10842442 | 1:200 |

**Table S2 – Secondary antibodies for IHC, relating to Fig.1-7, S1-3 and S5-6.**

| | Manufacturer | Product code | RRID | Dilution |
|---|---|---|---|---|
| 488 Donkey anti-rabbit | ThermoFisher | A21206 | AB_2535792 | 1:400 |
| 488 Donkey anti-goat | ThermoFisher | A11055 | AB_2534102 | 1:400 |
| 488 Donkey anti-mouse | ThermoFisher | A21202 | AB_141607 | 1:400 |
| 546 Donkey anti-rabbit | ThermoFisher | A10040 | AB_2534016 | 1:400 |
| 546 Donkey anti-goat | ThermoFisher | A11056 | AB_2534103 | 1:400 |
| 647 Donkey anti-rabbit | ThermoFisher | A31573 | AB_2536183 | 1:400 |
| 647 Donkey anti-goat | ThermoFisher | A21447 | AB_2535864 | 1:400 |
| 647 Donkey anti-mouse | ThermoFisher | A31571 | AB_162542 | 1:400 |
| 647 Donkey anti-chicken | ThermoFisher | A78952 | AB_2921074 | 1:400 |

**Table S3 – Primers used in this study for qRT-PCR analysis, related to Figs.5-7, S3 and S5.**

| Gene | Forward | Reverse | Universal Probe |
|---|---|---|---|
| *AIPL1* | GGAGAGGGAAATCGGCTCTT | TTCTCCTTGGTCTGCAGGTT | 79 |
| *ALDH1A1* | GTTAGCTGATGCCGACTTGG | CGCTCAACACTCCTTCGAAC | 81 |
| *ALDH1A3* | AAGAGGAGATTTTCGGGCCA | ACTTCAGGGCTTTGTCGAGA | 66 |
| *ARR3* | CACGGAGACTGTAGCTGCTA | GCTAGAGGCCAGGTTGGTAT | 14 |
| *β-ACTIN* | GCACCCAGCACAATGAAGAT | TCGTCATACTCCTGCTTGCT | 63 |
| *CRX* | GAGGCTGTGTCTTGTGCAAA | TCGAAGTCTTACCCCATCGG | 12 |
| *CRABP2* | CCCTGTAAGAGCCTGGTGAA | CCGTCATGGTCAGGATCAGT | 93 |
| *CYP26A1* | GACATGCAGGCACTAAAGCA | GGTAGAGCCCCAGGTAAGTG | 5 |
| *FGF8* | GAGACGGGCCTCTACATCTG | TGTACCAGCCCTCGTACTTG | 67 |
| *FGFR1* | CGGGACAGACTGGTCTTAGG | GGTTGGGTTTGTCCTTGTCC | 53 |
| *FGFR2* | CAGTCATCCTGTGCCGAATG | AACTGTTACCTGTCTCCGCA | 71 |
| *FGFR3* | TACTCCTTCGACACCTGCAA | ACTTCTGGGAGGCCAAGTAC | 68 |
| *FGFR4* | GGAGTCCCGGAAGTGTATCC | GTAGTCAATGTGGTGGACGC | 71 |
| *GAPDH* | TTGCCCTCAACGACCACTTT | TGGTCCAGGGGTCTTACTCC | 77 |
| *ISLET1* | TCTCCGGATTTGGAATGGCA | CATTTGATCCCGTACAACCTGAT | - |
| *L-OPSIN* | TGCATCATCCCACTCGCTAT | TTCCTTCTCTGCCTTCTGGG | 5 |
| *M-OPSIN* | CTACCTCCAAGTGTGGCTGG | GAATGCCAGGACCATCACCA | 13 |
| *M/L-OPSIN* | CTTCCACCCTTTGATGGCTG | AGTTTCGAAACTGCCGGTTC | - |
| *NRL* | AGAGCTGTGTGCAGAAGTCT | GCAGAAGTCGTCCAATCCAC | 95 |
| *ONECUT1* | CAGCGTCGAACTCTACATGC | ACTCCTCCTTCTTGCGTTCA | 3 |
| *OTX2* | ATTGCTAGAGCAGCCCTCAC | GGAGCAGTGGAACTTACAGC | - |
| *RBPMS* | CACTGCATGCCCAGATGC | CCCACAAGACAGATTGCAGC | - |
| *RECOVERIN* | GCTTCGATTCCAACCTCGAC | TTACCGTCCACGTCGTAGAG | 78 |
| *RHODOPSIN* | TGGGCCATTAAAAGCTCAGC | GTGGAAGCTGGCAGTTTCAA | 77 |
| *RXRγ* | CATGAAGAGGGAAGCTGTGC | TTCTGCCTCACTCTCAGCTC | 82 |
| *SFRP1* | GCCCGAGATGCTTAAGTGTG | GACACACCGTTGTGCCTTG | - |
| *S-OPSIN* | GCACTGTAGCAGGTCTGGTT | CAATGGTCCAGGTAGCCAGG | 72 |
| *THRβ* | ATTTCGTTCTCCGGTCCTGT | TCAAGTCTTGGACCAGGGAC | 9 |
| *VSX2* | TGGGGATGCACAAAAAGTCG | GCTCCATCTTGTCGAGCTTG | 69 |

### Supplementary Experimental Procedures

#### Human PSC culture and retinal differentiation

Established hESC line H9 (Wicell: WAe009-A, lot RB66492, passage (P)30) cells were maintained on Geltrex-coated (Gibco: A1413302) 6-well plates and fed daily with E8 media (Gibco, A1517001). Cells were passaged when 80% confluent in clumps using Versene (Gibco, 15040066) and plated 200,000 cells per well with 1:1000 Rock Inhibitor Y-27632 dihydrochloride (Tocris: 1254). Cells were cryopreserved with Knockout Serum Replacement (Thermo: 10828010) and 10% DMSO (Generon, DMSO-10) in liquid nitrogen.

Authentication of the Master Cell Bank by STR (Short Tandem Repeat) analysis was performed by WiCell, and confirmed exact match of the STR profile. Karyometrix testing was conducted on the working cell bank (P33), and the last culture passage (P64), with no major chromosomal changes detected at either stage. Pluripotency validation was performed for the working cell bank via flow cytometry of intracellular (Nanog, >70%; Oct3/4, Sox2; >80%) and extracellular markers (SSEA-1, <10%; SSEA-3, TRA-1-81, >80%) alongside positive immunocytochemistry staining for Nanog, Oct3/4 and Sox2. All experiments were performed on PSCs <15 passages after thawing from the working cell bank. During culture of both PSCs and hROs, regular mycoplasma and sterility testing was performed monthly.

For differentiation, human PSCs were maintained until 90-95% confluent, then on day 1 (d1) cultured in E6 media (Gibco, A1516401) for 2 days. Proneural induction media (PIM) was added for 40 days, changed every 2-3 days, consisting of Advanced DMEM/F12 (Gibco: 12634-010), N2 (Gibco 17502-048), Glutamine (Thermo Scientific:25030-024), Antibiotic-Antimycotic (Gibco: 15240-062), Non-essential amino acids (Gibco: 11140-035). Optic vesicles were manually excised from day 21 with 21G needles and cultured in low-binding 96-well plates in retinal differentiation media (RDM), consisting of: DMEM (Gibco: 41965-039), F12 (Gibco: 31765-027), B27-vit A (Gibco: 12587-010), Antibiotic-Antimycotic (Gibco: 15240-062). At day 42, additional factors were added to RDM: 10% FBS (Gibco: 10500-064), 2mM Glutamax (Gibco: 35050-038), 100uM Taurine (Sigma: T4571). From day 70, media was changed to Alt70, consisting of: Advanced DMEM/F12 (Gibco: 12634-010), B27-vit A (Gibco: 12587-010), Antibiotic-Antimycotic (Gibco: 15240-062), 10% FBS (Gibco: 10500-064), 2mM Glutamax (Gibco: 35050-038), 100uM Taurine (Sigma: T4571) with 1uM retinoic acid added fresh. From day 90 media was changed to Alt90, consisting of: Advanced DMEM/F12 (Gibco: 12634-010), B27-vit A (Gibco: 12587-010), N2 (Gibco 17502-048), Antibiotic-Antimycotic (Gibco: 15240-062), 2mM Glutamax (Gibco: 35050-038), 100uM Taurine (Sigma: T4571), with 0.5uM retinoic acid added fresh (Biotechne: 0695/50).

##### Fetal retina/hRO sample preparation for Immunohistochemistry/ RNAscope™

For immunohistochemistry (IHC)/RNAscope™, human fetal samples were fixed for 1 day in 10% formalin before transferring to PBS. Samples were incubated overnight in 20% (w/v) sucrose (Merck: 84100), prior to cryo-embedding in OCT matrix (CellPath: 15212776). Tissue was cut at 10mM thickness, and mounted on Superfrost glass slides (Epredia: 10149870). hROs were washed with PBS and fixed for 1 hour in 4% PFA (Merck: P6148), followed by PBS wash and incubated overnight in 20% (w/v) sucrose, prior to cryo-embedding in OCT matrix. Tissue was cut at 10mM thickness, and mounted on Superfrost glass slides, air-dried overnight and frozen at -20°C.

##### RNAscope™ *in situ* hybridization

Samples were baked at 60°C for 30 minutes, postfixed with 4% PFA for 15 minutes at 4°C then dehydrated using an ethanol gradient, before 10-minute treatment with hydrogen peroxidase. Manual antigen retrieval was performed at 80°C for 5 minutes, then sections were pretreated with protease III for 30 minutes at 40°C. Sections were incubated with probes from the RNAscope™ probe catalogue Hs-CYP26A1-C1 (ACD Bio: 487741) and Hs-FGF8-C2 (ACD Bio: 415791-C2), or Human 3-plex positive (ACD Bio: 320861 – *POLR2a*/*PPIB*/*UBC*)/negative controls (ACD Bio: 320871 – DapB) for 3 hours at 40°C, before storing in 5xSSC (NaCl: Merck: S9625; Sodium Citrate: Thermo: 045556.30) overnight. Probes went through series of amplification steps, according to the manufacturer's protocol, before developing the signal and assigning a fluorochrome to each probe by incubating for 30 minutes with PerkinElmer cyanine 3/5 fluorophores, respectively. Following HRP blocking, slides were counterstained with VSX2 by incubating with primary antibody overnight, and secondary antibody Alexa-fluor 488 (**Table S2**)/DAPI (Invitrogen: D1306) for 2 hours, according to the IHC protocol and mounted in DAKO mounting medium (Agilent: S302380-2). Confocal images were taken using a Leica DM5500Q and processed in ImageJ. All image adjustments, including brightness, were applied uniformly to both experimental and control images.

Mean fluorescence intensity of *CYP26A2*/*FGF8* in each sample image (PM/ONH) as well as positive and negative controls for the corresponding channels was calculated using ImageJ (**Fig.S1b**). Fluorescence intensity of negative control was subtracted, to control for background noise, then intensity was normalized to positive control, to account for technical variation between samples. One-way ANOVA and Tukey's multiple comparison tests were performed in GraphPad Prism (v10.1.2).

#### Immunohistochemistry (IHC)

Cryosections were permeabilized for 1 hour at room temperature with PBS/0.1% Tween (Merck: P9416)/0.5% Triton X (Merck: T8787), then blocked for 2 hours at room temperature with PBS/0.1%

Tween/5% donkey serum/1% BSA (Merck: A9418). Primary antibodies with incubated overnight at 4°C (**Table S1**). Alexa Fluor® secondary antibodies (**Table S2**) were incubated for 1 hour at room temperature at 1:400 dilution, along with DAPI counterstain. Confocal images were taken using a Leica DM5500Q and processed in ImageJ. All image adjustments, including brightness, were applied uniformly to both experimental and control images. Analysis was performed using unpaired t-tests. Spot count analysis was performed using Imaris 10.0. For analysis of KI67 in human fetal tissue, the same length of region was selected at different regions of the same retina: the PM (identified by *CYP26A1*+ staining); the central region between the PM and nerve; the region the same distance as the PM from the periphery on the nasal side of the retina; and the periphery (**Fig.S2a**). For analysis of CAPSASE staining in hROs, necrotic core was masked to ensure only cell death in the neural retina cells was included (**Fig.S6**).

##### qRT-PCR gene expression analysis

For all qPCR assays, 3-5 hROs were pooled for each sample depending on collection timepoint (5 for d40-d90, 3 for d120-150). Samples were taken from independent experiments and a minimum of n=3 independent samples were used for analysis. RNA was extracted from hRO pellets using RNeasy Micro kit (Qiagen: 74004), according to manufacturer's instructions. Samples were eluted in 30ml and stored at -80°C. Concentration and RNA quality analyzed using a BioDrop. cDNA was synthesized using QuantiTect Reverse Transcription Kit (Qiagen: 205311), according to manufacturer's instructions. Quantitative RT-PCR was performed using Perfecta Low-ROX MasterMix (Quantabio: 95120-012) with custom primers listed in **Table S3** and probes from the Merck Universal Probe library or PowerTrack SYBR MasterMix (Applied biosystems: A46111) with ROX as passive reference. All samples were run in triplicate alongside 2 endogenous reference genes (*GAPDH* and *b-actin*), along with water and no RT negative controls. Comparative CT was calculated to determine relative gene expression as fold change from either d0 PSCs or untreated controls. Treated/untreated samples were from the same batch of differentiation to control for batch-batch variation, and paired T-tests/one-way repeated measures ANOVA followed by Dunnett's test against control group was performed (adjusting for multiple comparisons).

#### hRO growth analysis

Growth of hROs was analyzed by measuring organoid area with ImageJ from brightfield images before and after dosing, from which % increase in area size calculated for each hRO.

#### Alamar Blue Viability assay

AlamarBlue utilizes resazurin which is reduced in presence of living cells and is non-toxic/cell permeable. Organoids were kept in separate wells of a 96-well plate and incubated for 4 hours with 10% AlamarBlue (Invitrogen: DAL1025) diluted in media, after which AlamarBlue was transferred to a new 96 well plate, organoids were washed 3 times with PBS and returned to media. Fluorescence of AlamarBlue at 590nm (RFU) was then measured using a plate scanner and % reduction calculated using:

$$Reduction_{AB} = \frac{RFU^{Exp} - RFU^{Neg}}{RFU^{100\%} - RFU^{Neg}} \times 100$$

In which $Reduction_{AB}$ is the percentage reduction of AlamarBlue, $RFU^{Exp}$ is the Relative Fluorescence Units (RFU) for the experimental sample, $RFU^{Neg}$ is the RFU for the Negative control, and $RFU^{100\%}$ is the RFU for the 100% reduced positive control.

To control for differences in hRO size, change in reduction from day 0 was calculated for each hRO:

$$\Delta Reduction = Reduction_{AB}^{Day\ X} - Reduction_{AB}^{Day\ 0}$$

Relative change in viability of treated organoids was normalized to DMSO treated controls:

$$\Delta Viability = \frac{\Delta Reduction^{Exp}}{\Delta Reduction^{Control}} \times 100$$

In which *ΔViability* is the change in viability from Day 0 in the experimental sample relative to the DMSO-dosed control sample

As hRO viability was significantly reduced upon dosing with 20µM AGN19309 but not 10µM, this concentration was used for all following experiments (**S4a Fig**). No reduction in viability was observed at any concentration of rFGF8 (**S4b Fig**). FGF8 can self-regulate its mRNA expression[56]. Consequently, expression of *FGF8* was assessed following dosing with rFGF8. No significant difference in *FGF8* expression was observed at any concentration, however, the largest mean difference was found at 100ng/µl (**S4b Fig**). Consequently, this concentration was used for all following experiments.

#### Statistics

All data are presented as mean±SD; N denotes number of independent experiments (i.e., differentiation batches) and n denotes number of images or hROs examined, where appropriate. Statistical testing was performed in GraphPad Prism (version 10.1.2). Tests used to analyze statistical significance are specified in methods/figure legends. Outliers were identified and excluded using the ROUT test (Q=10%). Shapiro-Wilk Tests were used to assess normality and F-tests were used to confirm equal variances prior to applying parametric statistical tests.
